# Self-restrained sex chromosome drive through sequential asymmetric meiosis

**DOI:** 10.1101/2024.11.14.623699

**Authors:** Xuefeng Meng, Yukiko M. Yamashita

## Abstract

Meiotic drivers are selfish genetic elements that bias their own transmission, violating Mendel’s Law of Equal Segregation. It has long been recognized that sex chromosome-linked drivers present a paradox: their success in transmission can severely distort populations’ sex ratio and lead to extinction. How sex chromosome drivers may resolve this paradox remains unknown. Here, we show that *D. melanogaster*’s *Stellate (Ste)* is an X chromosome-linked driver with a self-restraining mechanism that weakens its drive and prevents extinction. Ste protein asymmetrically segregates into Y-bearing cells during meiosis I, subsequently causing their death. Surprisingly, Ste segregates asymmetrically again during meiosis II, sparing half of the Y-bearing spermatids from Ste-induced defects. Our findings reveal a novel class of sex chromosome drivers that resolve the paradox of suicidal success.

## Main Text

Meiotic drive is a phenomenon in which a genetic element (the meiotic driver) is transmitted to offspring at a rate higher than predicted by Mendel’s Law of Equal Segregation (*1–4*). Meiotic drivers are proposed to selfishly propagate within the population, even at the cost of the host’s fitness, and may exert a strong evolutionary force (*2–4*). Many drive systems are linked to sex chromosomes, leading to the non-Mendelian transmission of one sex chromosome over the other in the heterogametic sex, thereby distorting the sex ratio in the progeny (*5*). However, sex chromosome-linked drivers present a paradox: if a driver is successful and strongly skews the sex ratio in offspring, it would eventually drive the population to extinction, thus preventing its own transmission (*6–8*). This paradox was theorized by W. D. Hamilton, who used mathematical modeling to demonstrate that a strong sex ratio distortion rapidly leads to population extinction (*9*). The mechanism by which sex chromosome drivers avoid this fate has remained a mystery.

*Stellate (Ste)*, an X chromosome-linked multicopy gene in *Drosophila melanogaster*, is normally repressed by piRNAs produced from the Y chromosome-linked *Suppressor of Stellate* (*Su(Ste)*, also known as *crystal* (*cry*)) (Fig. 1A) (*10–13*). *Ste* is a suspected meiotic driver that biases transmission of the X chromosome in males (*14, 15*), yet previous studies found only weak sex ratio distortion (60-80% female) upon *Ste* derepression. Additionally, a higher degree of *Ste* expression did not proportionally lead to increased transmission of the X chromosome, calling into question the identity of *Ste* as a meiotic driver (*16–18*). Here, we demonstrate that *Ste* is indeed a meiotic driver; however, it operates through a ‘self-restraining’ mechanism that weakens the drive strength. This mechanism facilitates the propagation of the *Ste-*encoding X chromosome within the population while preventing extinction caused by an extremely distorted sex ratio.

**Fig. 1.**
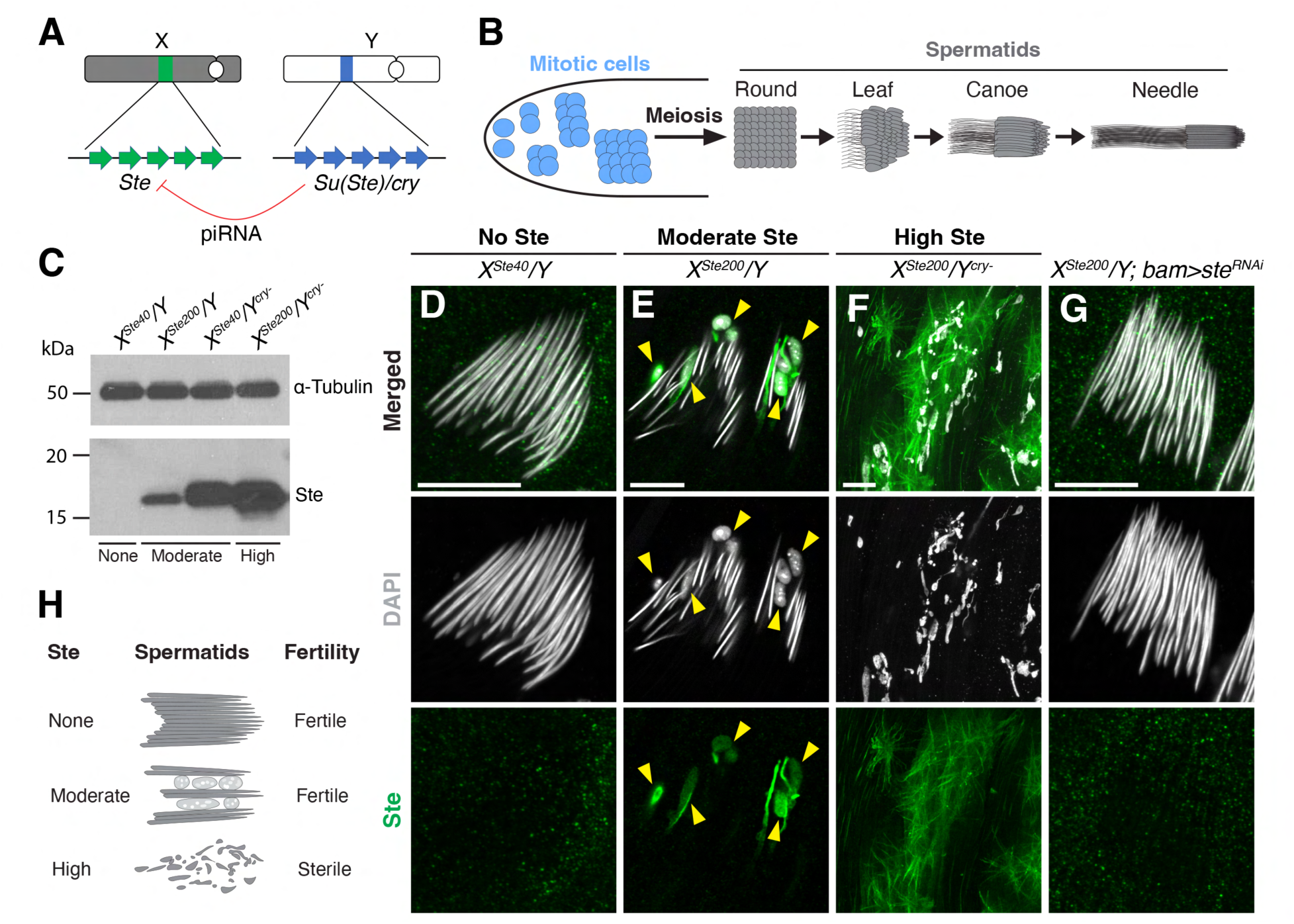
Moderately expressed Ste leads to defective sperm nuclear compaction. (**A**) Schematic showing the locations of *Ste* and *Su(Ste)/cry* on the X and Y chromosomes. *Ste* is typically silenced by piRNAs produced from *Su(Ste)*. (**B**) Schematic of *Drosophila melanogaster* male germ cell differentiation. (**C**) Western blotting of whole testis lysates from the indicated genotypes, probed with anti-α-Tubulin (loading control) and anti-Ste antibodies. (**D** to **G**) Immunofluorescence staining for Ste (green) in needle-stage spermatid cysts of *X^Ste40^/Y* (D), *X^Ste200^/Y* (E), *X^Ste200^/Y^cry-^* (F), and *X^Ste200^/Y; bam>ste^RNAi^* (G). Yellow arrowheads indicate sperm nuclei with DNA compaction defects. Grey, DAPI. Scale bars, 10 μm. (**H**) Diagram showing phenotypes under different *Ste*-expressing conditions.

### Ste impedes sperm nuclear DNA compaction

Previous studies have shown that *Ste* can be derepressed to varying degrees, depending on both *Ste* copy number and the activity of the piRNA pathway. In this study, similar to previous research (*16, 19*), we used combinations of methods to achieve varying levels of *Ste* derepression: (1) X chromosomes with different *Ste* copy numbers (X^Ste40^ with ∼40 copies and X^Ste200^ with ∼200 copies of *Ste*) (fig. S1A), (2) wild-type versus *Su(Ste)/cry*-deleted Y chromosomes (fig. S1B), and (3) knockdown of the piRNA pathway component Aubergine (Aub) (Fig. 1C, and fig. S1C, see table S1 for more detailed description). These manipulations resulted in conditions in which *Ste* was either not expressed, moderately expressed, or highly expressed. In conditions without Ste (i.e., wild-type or equivalent), no cytological phenotype was observed, thus serving as a control (Fig. 1D, and fig. S1D). In high Ste conditions, due to catastrophic meiosis (*16*), no functional post-meiotic spermatids were produced and the males were completely sterile (Fig. 1F, and fig. S1, H to J) (*16, 20, 21*). In moderate Ste conditions, *Ste* was not expressed at levels high enough to cause sterility (fig. S1J), but it led to sex ratio distortion skewed towards female progeny (60-80% female) as previously reported (*16, 18*) (see below for the sex ratio assay). In this study, we used multiple genotypes (*X^Ste200^/Y*, *X^Ste40^/Y*; *aub^RNAi^*, and *X^Ste40^/Y^cry-^*) that induce moderate *Ste* expression (table S1), and we investigated how *Ste* leads to meiotic drive.

Consistent with earlier studies (*16, 18*), males with moderate *Ste* expression remained fertile (fig. S1J). Under this condition, spermatogenesis appeared mostly normal (fig. S1, E to G), with all stages of differentiating germ cells present in an apparently normal spatiotemporal order within the testis. However, abnormalities became apparent during post-meiotic spermatid development. In wild-type flies, post-meiotic sperm differentiation was accompanied by stereotypical morphological changes, resulting in highly compacted sperm nuclei (Fig. 1, B and D). In flies with moderate *Ste* expression, we observed Ste protein localizing to a subset of spermatids throughout the post-meiotic stages (fig. S2, B to E, G to J), which eventually exhibited nuclear DNA compaction defects (Fig. 1E, and fig. S2, E and J). RNAi-mediated knockdown of Ste rescued the nuclear DNA compaction defect (Fig. 1G, and fig. S1K), confirming that this defect was indeed caused by *Ste* expression. The observed nuclear DNA compaction defects resemble those induced by other known ‘sperm-killing’ meiotic drivers, such as *D. melanogaster Segregation Distorter (SD)* (*22, 23*) and *D. simulans Sex Ratio (SR)* (*24*). Similar to *SD* (*23*), Ste-containing spermatids that failed to compact DNA also failed to incorporate protamines, such as Mst77F and ProtB, which are essential components of sperm chromatin (*25–27*) (fig. S3, B to D). These defective nuclei also lacked histones, indicating that histones were removed without undergoing the proper histone-to-protamine transition (*28*) (fig. S3, A and B). Taken together, we conclude that moderately expressed Ste protein localizes to a subset of differentiating spermatids, leading to defective sperm development and ultimately sperm death (Fig. 1H).

### Ste preferentially harms Y-bearing sperm

Moderate *Ste* expression is known to mildly increase the proportion of females in the progeny, which originally led to the hypothesis that *Ste* is a meiotic driver (*14, 15*). To test whether Ste-mediated defects in sperm development preferentially affect Y-bearing sperm, we performed DNA FISH using probes specific to the X or Y chromosome ((TAGA)_n_ for the X and (AATAAAC)_n_ for the Y) (*29*). We found that the majority of Ste-containing spermatids carried Y chromosomes (Fig. 2, A to C, and fig. S4, A and B), suggesting that Ste acts as a meiotic driver by preferentially killing Y-bearing spermatids. Moreover, we found that transgenic piRNA-resistant Ste (*β-tubulin promoter-Ste^piRNA-resistant^*) also preferentially localized to Y chromosome-bearing spermatids and caused nuclear DNA compaction defects (fig. S5). Together with the finding that RNAi-mediated knockdown of Ste rescues the nuclear DNA compaction defect (Fig. 1G, and fig. S1K), these results demonstrate that Ste is both necessary and sufficient to cause the biased killing of Y-bearing sperm.

**Fig. 2.**
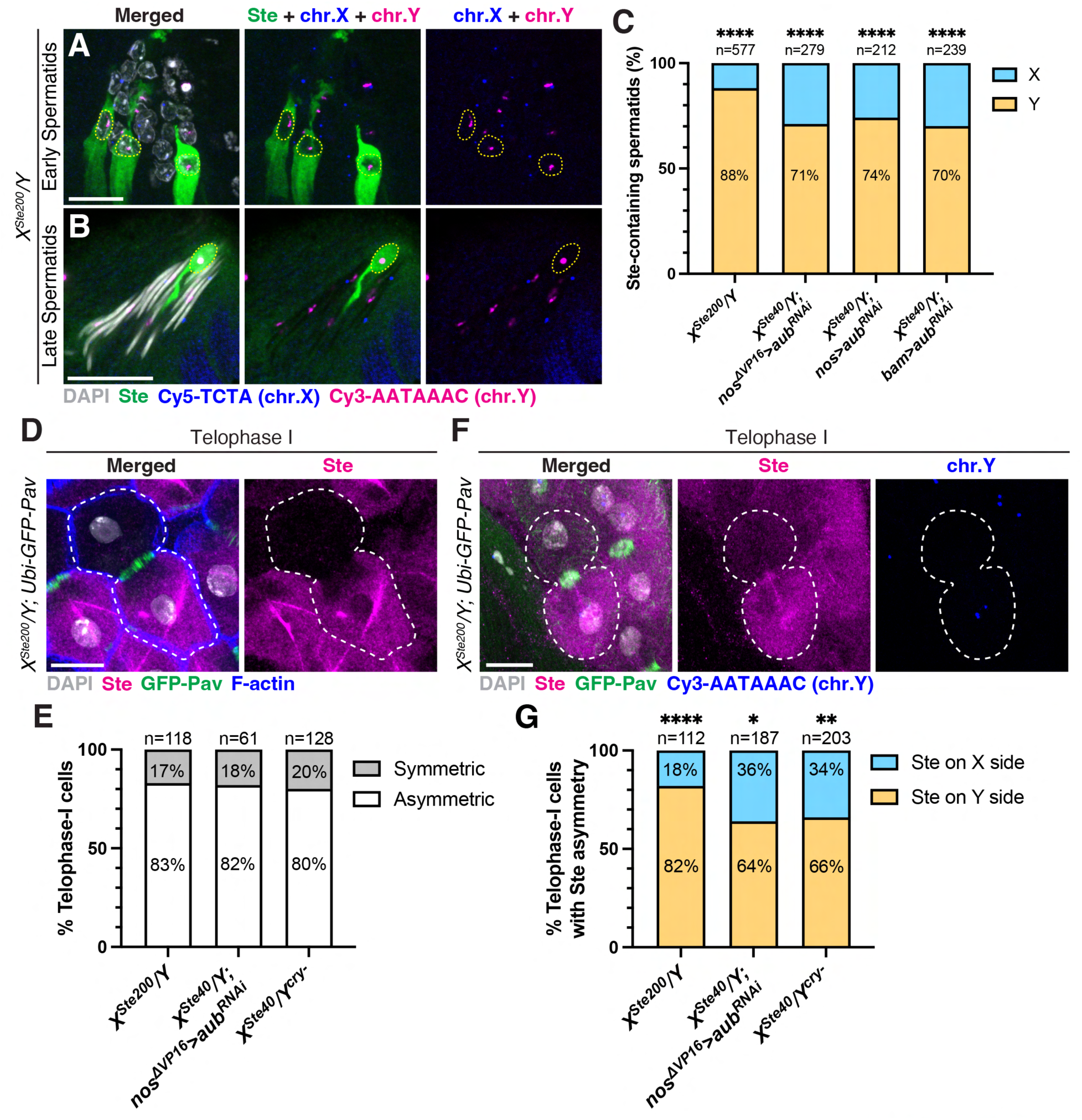
Ste preferentially segregates to Y-bearing spermatids through asymmetric segregation during meiosis I. (**A** and **B**) Immunofluorescence staining for Ste (green) combined with DNA-FISH for X- and Y-chromosome-specific satellite DNA sequences (X: Cy5-TCTA, blue; Y: Cy3-AATAAAC, magenta) in early (A) and late (B) spermatid cysts of *X^Ste200^/Y* males. Nuclei of Ste-containing spermatids are indicated by yellow dotted lines. Grey, DAPI. Scale bars, 10 μm. (**C**) Percentage of X- or Y-bearing spermatids among Ste-containing spermatids in the indicated genotypes. The number of scored Ste-containing spermatids is shown above the bar graph. Statistical analysis was performed using two-sided Fisher’s exact tests (Null hypothesis: Ste-containing spermatids have equal chances of carrying the X or Y chromosome). \*\*\*\**P* < 0.0001. (**D**) Immunofluorescence staining for Ste (magenta) combined with Phalloidin staining (F-actin, blue) in a telophase I cell (indicated by white dotted lines) of *X^Ste200^/Y*; *Ubi-GFP-Pav* telophase I cells displaying asymmetric segregation of Ste protein in the indicated genotypes. The number of scored telophase I cells is shown above the bar graph. (**F**) Immunofluorescence staining for Ste (magenta) and GFP-Pav (green), combined with DNA-FISH for the Y chromosome-specific satellite DNA sequence (Cy3-AATAAAC, blue) in a telophase I cell (indicated by white dotted lines) of *X^Ste200^/Y*; *Ubi-GFP-Pav* males. Grey, DAPI. Scale bar, 10 μm. (**G**) Percentage of telophase I cells with Ste co-segregating with the X or Y chromosome among cells with asymmetric Ste segregation in the indicated genotypes. The number of scored telophase I cells is shown above the bar graph. Statistical analysis was performed using two-sided Fisher’s exact tests (Null hypothesis: Ste has equal chances of co-segregating with the X or Y chromosome during meiosis I). \**P* = 0.0121, \*\**P* = 0.0018, \*\*\*\**P* < 0.0001.

### Ste asymmetrically segregates with the Y chromosome in meiosis I

We next investigated how the Ste protein becomes concentrated in Y-bearing spermatids. Ste’s preferential localization in Y-bearing spermatids was evident immediately following meiosis (Fig. 2A, and fig. S4A) and persisted throughout spermatid development (Fig. 2, A and B, and fig. S4, A to C). In contrast, Ste was expressed in all spermatocytes immediately before meiosis (fig. S2, A and F). These results suggest that Ste’s preferential localization to Y-bearing spermatids is established during meiosis. Indeed, we observed that Ste protein was asymmetrically enriched in only one daughter cell in over 80% of meiotic telophase I cells (Fig. 2, D and E, and fig. S6, A and C). Strikingly, in the majority of the cases, Ste and the Y chromosome co-segregated to the same side during telophase I (Fig. 2, F and G, and fig. S6, B and D). Although Ste protein is known to form amyloid-like aggregates (crystals) (*20, 30, 31*), we also noted a population of diffusely distributed Ste protein within the cell, which was also asymmetrically enriched in the same side as the Ste aggregates in telophase I cells (Fig. 2, D and F, and fig. S6). Based on these observations, we conclude that Ste preferentially localizes to Y-bearing spermatids because of its asymmetric segregation during meiosis I.

### Ste segregates asymmetrically in meiosis II

Our findings thus far support that *Ste* is a meiotic driver that biases transmission of the X chromosome to offspring by harming sperm DNA compaction of Y-bearing spermatids. However, questions remain unanswered as to why Ste causes only a mildly increased female-to-male ratio, and why the frequency of X-bearing sperm (i.e., female offspring) does not increase proportionally with increasing amount of Ste, as noted decades ago (*16–18*).

To our surprise, we discovered that Ste undergoes asymmetric segregation during meiosis II as well (Fig. 3, A and B, and fig. S7, A and C). During telophase of meiosis II, approximately 80% of Ste-containing cells exhibited asymmetric segregation of Ste (Fig. 3B). Thus, even if a cell inherited Ste protein at the end of meiosis I, meiosis II would result in one spermatid with Ste and the other without it (Fig. 3D). We noted that the asymmetric segregation pattern during meiosis II was independent of the sex chromosomes: whether cells were segregating X-X or Y-Y sister chromatids, Ste was asymmetrically segregated in ∼80% of cases (fig. S7E).

**Fig. 3.**
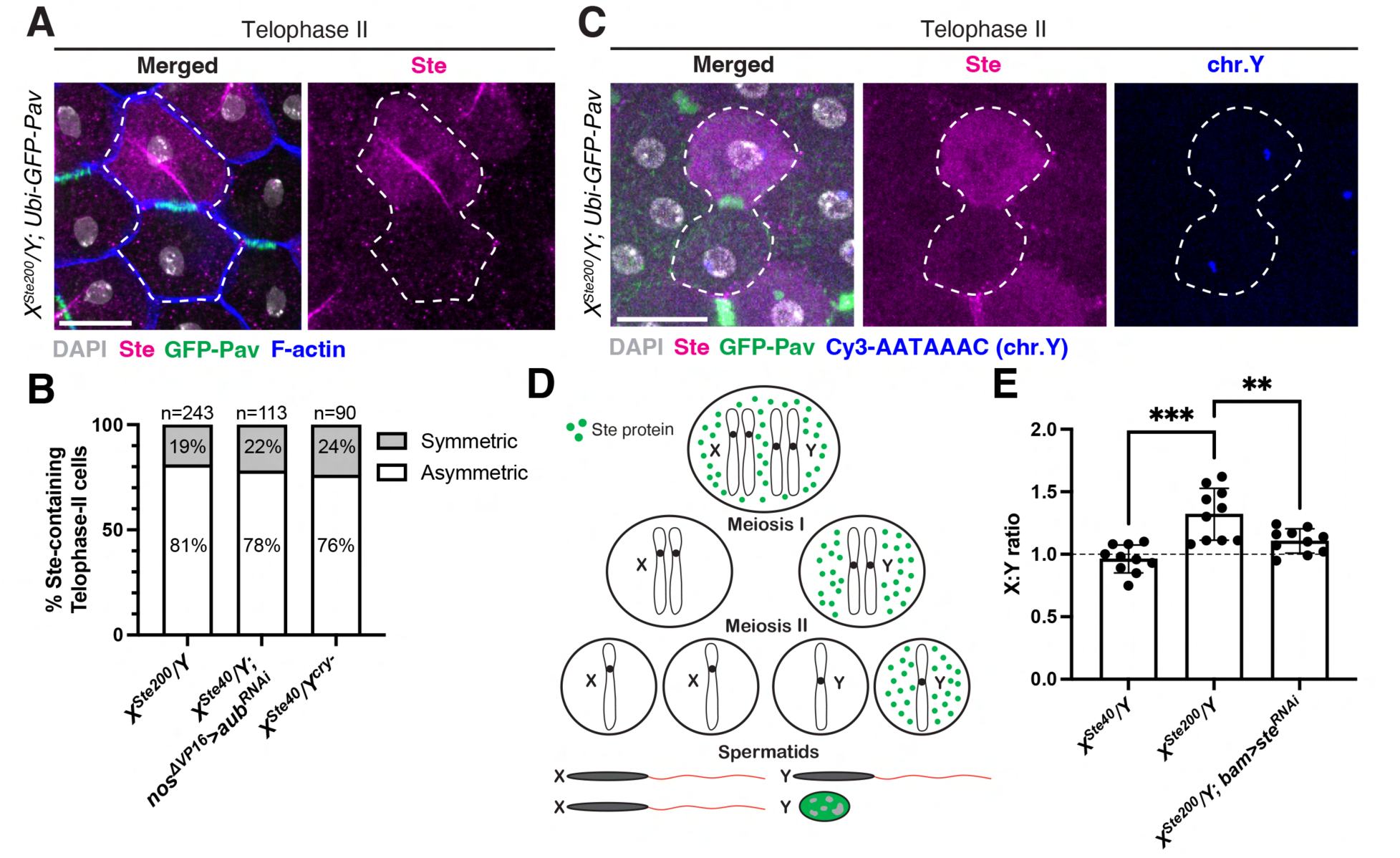
Ste exhibits asymmetric segregation during meiosis II. (**A**) Immunofluorescence staining for Ste (magenta) combined with Phalloidin staining (F-actin, blue) in a telophase II cell (indicated by white dotted lines) of *X^Ste200^/Y*; *Ubi-GFP-Pav* males (GFP-Pav, green). Grey, DAPI. Scale bar, 10 µm. (**B**) Percentage of Ste-containing telophase II cells displaying asymmetric segregation of Ste protein in the indicated genotypes. The number of scored telophase II cells is shown above the bar graph. (**C**) Immunofluorescence staining for Ste (magenta) and GFP-Pav (green), combined with DNA-FISH for the Y chromosome-specific satellite DNA sequence (Cy3-AATAAAC, blue) in a telophase II cell (indicated by white dotted lines) of *X^Ste200^/Y*; *Ubi-GFP-Pav* males. Grey, DAPI. Scale bar, 10 µm. (**D**) Model of asymmetric segregation of Ste during meiosis I and II, producing two X-bearing sperm and one Y-bearing sperm. Note that the frequency of asymmetry is not 100% and this model represents the scenario with the highest probability. (**E**) The ratio of X- to Y-bearing sperm produced by males of the indicated genotypes (calculated by the ratio of female to male progeny). Each dot in the graph represents a single male. Ten males were assayed for each genotype. Data are mean ± SD. Dashed line indicates the expected 1:1 ratio. Statistical analysis was performed using two-sided unpaired *t*-tests. \*\**P* = 0.0086, \*\*\**P* = 0.0001.

This asymmetric segregation during meiosis II has a striking implication: while Ste protein is preferentially segregated with the Y chromosome during meiosis I, about half of the Y-bearing spermatids are spared from inheriting Ste due to the asymmetric segregation in meiosis II (Fig. 3C, and fig. S7, B and D), allowing for the survival of roughly half of the Y-bearing spermatids (Fig. 3D). This explains why earlier studies observed only a mild female-biased sex ratio and why the female frequency did not increase proportionally with higher levels of *Ste* expression (*16–18*). We recapitulated this mild female-biased sex ratio in the progeny of *X^Ste200^/Y* males (Fig. 3E). Importantly, RNAi-mediated depletion of Ste rescued the sex ratio distortion (Fig. 3E), confirming that Ste is responsible for the skewed sex ratio. In conclusion, we demonstrate that Ste is indeed a meiotic driver; however, its asymmetric segregation during meiosis II results in only a weakly skewed sex ratio.

### Weak drive avoids extinction

It has long been recognized that a sex chromosome driver runs a risk of population extinction by skewing the sex ratio (*6–8*). Through mathematical modeling, Hamilton demonstrated that complete sex chromosome drive (100% transmission of the driving chromosome) would result in population extinction due to a severely skewed sex ratio (*9*). Our results described thus far suggest that the asymmetric segregation of Ste during meiosis II may serve as a mechanism to prevent *Ste* from becoming a complete X chromosome driver, which would lead to population extinction due to the depletion of Y chromosomes.

To explore how weak X chromosome drivers affect population dynamics, we conducted mathematical modeling. Using parameters similar to Hamilton’s model, we first recapitulated his results for 100% drive strength (complete drive) (Fig. 4, A and C): this strong drive caused rapid population extinction after a brief period of expansion, as shown previously (*9*). Interestingly, modulation of the drive strength (defined as the frequency of X-sperm produced by males, see Materials and Methods) significantly influenced population outcomes. Drive strengths of 90% or 80% also led to extinction, but at a slower rate than the 100% drive (Fig. 4A). In contrast, we found that lower drive strengths (70%, 60%) did not cause extinction, instead allowing continued population expansion (Fig. 4A). We found that there is a threshold drive strength that determines whether the population will eventually face extinction or not (Fig. 4B). When other parameters were kept consistent with Hamilton’s model (Materials and Methods), this threshold was 75% (Fig. 4B). Drivers stronger than 75% would shift from the positive growth phase (population change rate > 1) to the negative growth phase (population change rate < 1) after certain generations, ultimately leading to extinction (Fig. 4, B and C). Drivers with a strength of exactly 75% would reach a steady state (population change rate = 1) (Fig. 4, B and D). Any drivers with a strength between 50% and 75% would remain in the positive growth phase (Fig. 4B), resulting in continued population expansion (Fig. 4E). In contrast to a strong driver (strength = 100%) that rapidly leads to an extremely low male/female ratio in the population (Fig. 4C), weaker drivers that avoid extinction (strength = 75%, 60%) eventually reach a stabilized and less skewed sex ratio after many generations (Fig. 4, D and E). Notably, however, weak drivers do not compromise their ability to eventually fix themselves as the sole X chromosome in the population, although they do so at a slower rate than strong drivers (fig. S8A).

**Fig. 4.**
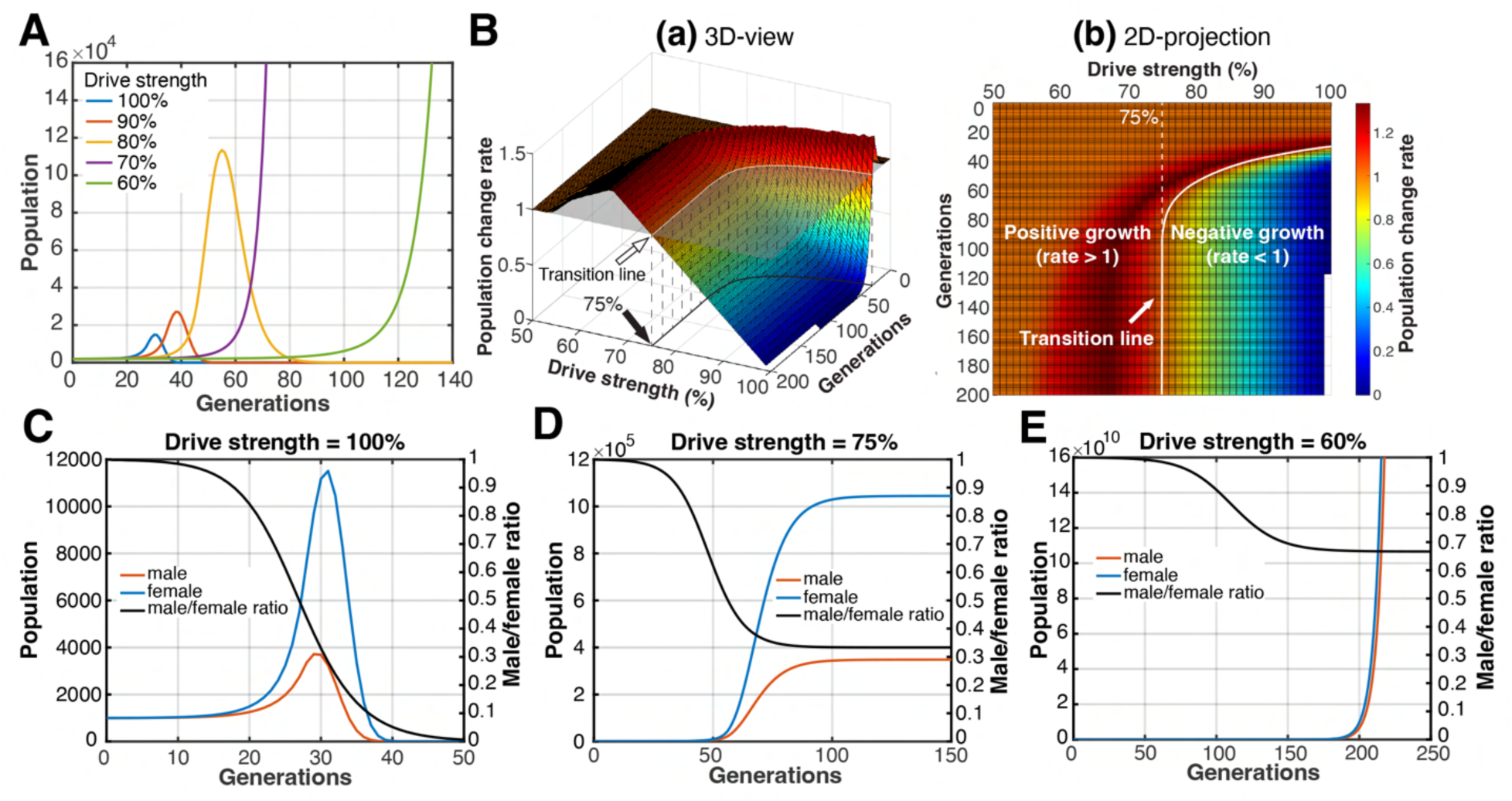
Weak sex chromosome drive avoids population extinction. (**A**) Population size across generations with varying degrees of drive strength (defined as the frequency of X-sperm produced by males). (**B**) (a) Population change rate over generations with varying degrees of drive strength. The transition line, where the population change rate is one (no population growth), is indicated by the white arrow. (b) Two-dimensional projection of (a), showing that the transition line separates the positive growth phase (rate > 1) from the negative growth phase (rate < 1). A driver with a strength of 75% will reach a steady state (no population growth) after a certain number of generations. (**C** to **E**) Simulation showing the male/female ratio (black line), number of females (blue line), and number of males (red line) across generations when the drive strength is 100% (C), 75% (D), and 60% (E).

While the exact threshold for drive strength is influenced by parameters such as the number of females a male can fertilize and the number of offspring a female can produce, changing these parameters still resulted in the presence of the threshold, which ranged >75% (fig. S8B, and Materials and Methods). Thus, drivers with a strength below 75% reside in a ‘safe zone’ to avoid eventual extinction. Interestingly, the observed drive strength for *Ste* (the frequency of X-sperm produced by *Ste*-expressing males) in this study and previous work (*16*) ranges from 56% to 82% with an average of 73% (fig. S8C, and table S2). These results indicate that *Ste*’s drive strength is likely below the threshold, allowing it to avoid extinction and resolving the paradox of sex chromosome meiotic drive.

## Discussion

Sex chromosome meiotic drive is believed to be an important evolutionary force (*2, 5*); however, it runs the risk of extinction due to an extremely skewed sex ratio that interferes with successful breeding (*6, 7, 9*). We propose that *Stellate* represents a novel class of meiotic drivers with a built-in mechanism to avoid the fate of extinction. The preferential segregation of Ste protein to the Y chromosome-bearing side during meiosis I provides the foundation for drive, while the asymmetric segregation during meiosis II serves as the mechanism to dampen the strength of drive. Our mathematical modeling shows that extinction is not the inevitable outcome of any X chromosome-linked drivers; only those exceeding a certain strength threshold lead to extinction. We propose that the asymmetric segregation of Ste during meiosis II weakens the drive strength below this critical threshold, allowing *Ste* to avoid the fate of extinction of a strong driver and resolving the paradox of sex chromosome drivers. It is worth noting that, while the rise of suppressors at distinct genetic loci is generally thought to counteract meiotic drivers (*3, 4*), *Ste* employs a self-restraining mechanism, making the driver inherently weak without the need for a suppressor, which could be separated from the driver through meiosis. This is not to say that the suppressing mechanism, i.e. *Su(Ste)*, is unnecessary. Fisher’s principle predicts a natural selection for a more balanced (1:1) sex ratio (*9, 32*), suggesting that *Su(Ste)* would be selected during evolution to counteract the distorted sex ratio caused by *Ste*. Additionally, the asymmetric segregation of Ste during meiosis II cannot prevent the meiotic failure caused by the high-level expression of *Ste*, thus requiring the action of *Su(Ste)*.

Meiotic drivers in females often utilize the inherent asymmetry of female meiosis by preferentially segregating into the egg and avoiding the non-transmissive polar bodies (gonotaxis), thus ensuring their transmission to the offspring (*3, 33–35*). In contrast, in males and fungi, where meiosis symmetrically produces four viable gametes from a diploid germ cell, meiotic drivers are often assumed to operate during the post-meiotic stage by sperm- or spore-killing (*3, 4*). This is because cytological defects typically appear in the post-meiotic stages (*22–24, 36–38*). However, it has been speculated that post-meiotic cytological defects may be caused by earlier events (*4, 39, 40*). Our study provides an example where sperm-killing can be seeded during meiosis, through the asymmetric segregation of a driver-encoded protein, which later kills sperm in the post-meiotic stages. It remains unclear whether the asymmetric segregation of Ste during meiosis is an implication of any unknown inherent asymmetry of male meiosis, similar to female meiosis. It also remains elusive how Ste co-segregates with the Y chromosome during meiosis I, which is the foundation of its drive. While Ste might have an affinity for the Y chromosome, the fact that Ste segregates asymmetrically during meiosis II, when identical sister chromatids are segregating away from each other, suggests that Ste’s asymmetry may not be entirely chromosome dependent.

In summary, we propose that *Ste* is a novel sex chromosome-linked meiotic driver that propagates without the risk of extinction through a built-in mechanism that self-restrains the strength of its drive. This type of ‘weak-by-design’ driver may have been overlooked and understudied due to its incomplete drive.

## Acknowledgments

We thank the members of the Yamashita lab for discussions and comments on the manuscript and help with experiments. We thank Drs. Ruth Lehmann, Ankur Jain, Scott Hawley, Andrew Clark, David Page, Gerald Fink, Jacob Mueller, and Christina Lilliehook for insightful comments and suggestions. We thank former Yamashita lab members Zsolt G. Venkei, Seoyeon (Chloe) Choi, Jonathan O. Nelson, George J. Watase, Madhav Jagannathan, Jullien Flynn, and Jun Park for insightful suggestions and help with experiments. We thank Peiwei Chen and Alexei Aravin for providing the sequence of the *βTub* promoter. We thank the Lehmann lab members for discussing the project. We thank the Flybase, Bloomington Stock Center, Developmental Studies Hybridoma Bank, and Kyoto Stock Center for reagents and critical information. We acknowledge ChatGPT (https://chat.openai.com/) for assistance with writing the code for mathematical modeling and for copy editing this manuscript.

## Funding

Howard Hughes Medical Institute (Y.M.Y)

## Author contributions

Conceptualization: XM, YY

Methodology: XM, YY

Investigation: XM

Visualization: XM

Funding acquisition: YY

Project administration: XM, YY

Supervision: YY

Writing – original draft: XM, YY

Writing – review & editing: XM, YY

## Competing interests

Authors declare that they have no competing interests.

## Data and materials availability

All data are available in the main text or the supplementary materials. MATLAB code for mathematical modeling is deposited at https://github.com/xuefengmeng/Meng_et_al_2024.git.

## Materials and Methods

### Fly husbandry and strains used

All *Drosophila melanogaster* strains were raised on standard Bloomington medium at 25 °C. The following strains were used: the standard lab wild-type strain *y w* (*y^1^w^1^*, used as *X^Ste40^*), the double balancer strain *X^Ste200^ /Y; Sp/CyO;TM2/TM6B*, *Y^cry-^* (*B^S^cry^1^Yy^+^*, a gift from Maria Pia Bozzetti) (*16, 18*), *nos-gal4^ΔVP16^* (*41*), *nos-gal4:VP16* (*42*), *bam-gal4:VP16* (Bloomington Drosophila Stock Center [BDSC]:80579, a gift from Dennis McKearin) (*43*), *ste^TRiP.HMJ30118^*(BDSC: 63552) (*44*), *aub^TRiP.GL00076^*(BDSC: 35201) (*44*), *Ubi-GFP-Pav* (BDSC: 81650, a gift from David Glover) (*45*), *Mst77F-EGFP* (Kyoto Stock Center DGRC: 109174) (*26*), and *ProtB-EGFP* (BDSC: 58406) (*26*).

### *βTub-Ste* transgene construction

The *βTub-Ste^piRNA-resistant^* transgenic strain was generated via phiC31 site-directed integration into the *Drosophila melanogaster* genome. The piRNA-resistant *Ste* cDNA was designed by introducing silent mutations throughout the entire CDS (sequence provided in table S3). We designed the CDS by adopting the consensus sequence from the 13 annotated *Ste* genes on FlyBase (Ste:CG33236, Ste:CG33237, Ste:CG33238, Ste:CG33239, Ste:CG33240, Ste:CG33241, Ste:CG33242, Ste:CG33243, Ste:CG33244, Ste:CG33245, Ste:CG33246, Ste:CG33247, and SteXh:CG42398). The piRNA-resistant cDNA was synthetized by Thermo Fisher Scientific (GeneArt Gene Synthesis) and inserted into the *pattB* vector along with the *β2-tubulin (βTub)* promoter (generously provided by Peiwei Chen and Alexei Aravin). The *pattB-βTub-Ste^piRNA-resistant^* construct was inserted into the *attP18* integration site on the X chromosome. The transgenic line was generated by BestGene Inc.

### Western blots

Testes (25 pairs per sample) from 2- to 3-day-old males were dissected, rinsed with 0.1 M phosphate buffer saline (1x PBS) at pH 7.2, snap-frozen, and stored at −80 °C until use. The testes were homogenized in 150 μl of 1x PBS supplemented with c0mplete protease inhibitor (EDTA-free, Roche) and mixed with 150 μl of 2× Laemmli Sample Buffer (Bio-Rad), supplemented with 2-mercaptoethanol. Cleared lysates were denatured at 95 °C for 3-5 minutes, separated on a 14% Tris-glycine gel (Thermo Fisher Scientific), and transferred onto a polyvinylidene fluoride membrane (iBlot2, Invitrogen). Mouse anti-α-Tubulin (AA 4.3; 1:3,000, obtained from the Developmental Studies Hybridoma Bank) and guinea pig anti-Ste (*46*) (used at 1:10,000) were used as primary antibodies. Horseradish peroxidase-conjugated goat anti-mouse IgG (115-035-003; 1:10,000; Jackson ImmunoResearch Laboratories) and anti-guinea pig IgG (106-035-003; 1:10,000; Jackson ImmunoResearch Laboratories) were used as secondary antibodies. Signals were detected using the Pierce ECL Western Blotting Substrate enhanced chemiluminescence system (Thermo Fisher Scientific).

### Immunofluorescence staining

Testes from 0- to 3-day-old males were dissected in 1x PBS and fixed in 4% formaldehyde in 1x PBS for 30 minutes. Fixed testes were then washed in 1x PBST (PBS containing 0.1% Triton X-100) for at least 2 hours, followed by incubation with primary antibodies diluted in 1x PBST containing 3% BSA at 4 °C overnight. Samples were washed three times in 1x PBST for 30 minutes each and then incubated with secondary antibodies in 1x PBST with 3% BSA at 4 °C overnight. After a similar washing procedure, samples were mounted in VECTASHIELD with DAPI (Vector Labs). Images were acquired using a Leica Stellaris 8 confocal microscope with a 63x oil immersion objective lens (numerical aperture 1.4) and processed with Fiji (ImageJ) software. The primary antibodies used were: anti-Ste (1:200; guinea pig) (*46*), anti-GFP (1:100; rabbit; Abcam, ab290), anti-histone H3 (1:200; rabbit; Abcam, ab1791), anti-Mst77F (1:100; guinea pig) (*47*), anti-Pavarotti (Pav) (1:100; rabbit; a gift from David Glover) (*48*), and anti-ATP5a (1:1000; mouse; Abcam, ab14748). Phalloidin-Alexa Fluor 568 (1:200; Thermo Fisher Scientific, A12380) was used to stain F-actin. Alexa Fluor-conjugated secondary antibodies (Life Technologies) were used at a 1:200 dilution.

### DNA fluorescence in situ hybridization

Testes from 0- to 3-day-old males were dissected as described above, and an optional immunofluorescence staining protocol (modified by adding 1 mM EDTA to formaldehyde, PBST, and BSA) was performed first when necessary. Subsequently, samples were post-fixed with 4% formaldehyde + 1 mM EDTA for 30 minutes and washed in 1x PBST + 1 mM EDTA for 30 minutes. Fixed samples were incubated with 2 mg/ml RNase A solution (in PBST) at 37 °C for 10 minutes, followed by washing with 1x PBST + 1 mM EDTA for 5-10 minutes. Samples were then rinsed in 2x SSC + 1 mM EDTA + 0.1% Tween-20 and washed in 2x SSC + 0.1% Tween-20 with increasing formamide concentrations (20%, 40%, and 50%) for 15 minutes each, followed by a final 30-minute wash in 2x SSC + 0.1% Tween-20 + 50% formamide. Hybridization mix (50% formamide, 10% dextran sulfate, 2x SSC, 1 mM EDTA, 1 mM probe) was added to the washed samples. Samples were denatured at 91 °C for 2 minutes and then incubated overnight at 37 °C. After hybridization, samples were washed three times in 2x SSC + 1 mM EDTA + 0.1% Tween-20 for 20 minutes each and mounted in VECTASHIELD with DAPI (Vector Labs). The following satellite DNA probes were used: Cy5-(TCTA)_8_ and Cy5-359 for the X chromosome, and Cy3-(AATAAAC)_6_ for the Y chromosome (*29*). The sequence of the Cy5-359 probe is AGGATTTAGGGAAATTAATTTTTGGATCAATTTTCGCATTTTTTGTAAG.

### Droplet digital PCR (ddPCR)

Genomic DNA was extracted from individual 0- to 3-day-old males and virgin females using a modified protocol of the DNeasy Blood and Tissue DNA extraction kit (Qiagen). Briefly, individual flies were homogenized in 200 μL of Buffer ATL containing proteinase K using a pipette tip in PCR tubes, followed by vortexing for 15 seconds and incubation at 56 °C for 1.5 hours. The samples were then transferred to 1.5 mL Eppendorf tubes and processed according to the manufacturer’s instructions. DNA samples were then quantified and checked for purity using a NanoDrop One spectrophotometer (Thermo Fisher Scientific). For ddPCR, 30 ng of DNA was used per 20 μL reaction for control genes (RpL and Upf1), 3 ng for *Ste*, and 0.3 ng for *Su(Ste)*. The primers and probes for control reactions were as described in our previous studies (*49, 50*), while those for *Ste* and *Su(Ste)* were designed by Bio-Rad. The specificities of *Ste* and *Su(Ste)* primers and probes were validated in fig. S1, A and B. ddPCR reactions were prepared according to the manufacturer’s protocol (Bio-Rad). In short, master mixes containing ddPCR Supermix for Probes (No dUTP) (Bio-Rad), DNA samples, and primer/probe mixes were assembled in PCR tubes and incubated at room temperature for 15 minutes to allow for restriction enzyme digestion. For *Ste* and *Su(Ste)* ddPCR, the HaeIII restriction enzyme (New England Biolabs) was used to digest repetitive DNA into single units. ddPCR droplets were generated from samples using the QX200 Droplet Generator (Bio-Rad) and underwent complete PCR cycling on a C100 deep-well thermocycler (Bio-Rad). Droplet fluorescence was read using the QX200 Droplet Reader (Bio-Rad). Sample copy numbers were determined using Quantasoft software (Bio-Rad). *Ste* and *Su(Ste)* copy numbers were calculated based on the copy numbers of reference genes RpL and Upf1. The copy number values determined by each control gene was averaged to determine the final copy number for each sample. Six flies were analyzed per genotype.

### Fertility and sex ratio assay

For each genotype, 10 males were assayed as follows. Individual 0- to 1-day-old males were crossed with two 1- to 4-day-old virgin *y w* females for 5 days at 25 °C and all F1 progenies were counted for fig. S1J. The raw data for the fertility assay are provided in table S4. For the sex ratio assay, individual 0- to 1-day-old males were crossed with two 1- to 4-day-old virgin *y w* females in a vial for 5 days at 25 °C. After 5 days of mating, females were discarded, and the same males were transferred to new vials to mate with two new 1- to 4-day-old virgin *y w* females for another 5 days at 25 °C. This mating scheme was repeated five times. The number and sex of progenies from each cross were recorded. The total female and male progenies from all five mating periods were used to calculate the X : Y (female : male) ratio for each male in Fig. 3E (table S5). The percentage of X-bearing sperm for each genotype was calculated based on the total number of female and male progenies from all 10 assayed males and is presented in fig. S8C, along with sex ratio data from previous study (*16*) (table S2). Notably, we observed an increase in the female/male progeny ratio as the male aged.

### Statistics and reproducibility

No statistical method was used to predetermine sample size due to the ample sample sizes afforded by the use of *Drosophila*. No data were excluded from the analyses. Data analysis was performed using Microsoft Excel and GraphPad Prism 10. All graphs, except those in Fig. 4 and fig. S8, A and B, were generated using GraphPad Prism 10. The experiments were not randomized, because this study did not involve treatment or exposure of animals to any agents. The Investigators were not blinded for data collection and analyses because methods used for data acquisition (immunofluorescence, DNA FISH, western blotting, fly number quantification, and ddPCR) are not influenced by the experimenter’s knowledge of fly genotype and the analyses were performed with the same automated algorithms. Two-sided Fisher’s exact tests were used for analyzing Ste’s preferential localization in spermatids and Ste’s preferential segregation during meiosis I (table S7 and S8). Note that a low frequency (0-4%) of non-disjunction events (XY or O spermatids) was observed (table S6) and was excluded from statistical analyses. Two-sided unpaired *t*-tests were used to compare the sex ratios of progeny in Fig. 3E. The exact *P* values are provided in each figure legend.

### Mathematical modeling

Mathematical modeling in Fig. 4 and fig. S8, A and B was performed using MATLAB. Code is deposited at https://github.com/xuefengmeng/Meng_et_al_2024.git. We designate the driving X chromosome as X’, and normal sex chromosomes as X and Y. The following symbols are assigned for generation *n*: *Total*_*n*_ (total number of flies), *M*_*n*_ (number of males), *F*_*n*_ (number of females), *A*_*n*_ (number of X’Y males), *B*_*n*_ (number of XY males), *C*_*n*_ (number of X’X’ females), *D*_*n*_ (number of X’X females), *E*_*n*_ (number of XX females), *R*_*n*_ (frequency of X’ in males), and *K*_*n*_ (frequency of X’ in females). Drive strength *t* is defined as the frequency of X-bearing sperm produced by males, where 0.5 < *t* ≤ 1. Same as Hamilton’s modeling (*9*), we initiate the population with 1000 females and 1000 males, with the frequency of the driving X chromosome (X’) in both females and males set at 1/1000 (*M*_0_ = *F*_0_ = 1000, *R*_0_ = *K*_0_ = 1/1000).

To simulate the effect of the driving X’ chromosome on the population, we calculate population sizes in each generation using the following method. If flies in generation *n* produce *N* eggs, then the number of X’ eggs is *N* · *K*_*n*_ and the number of X eggs is *N* · (1 − *K*_*n*_). Given that the frequency of X’Y males is *R*_*n*_, the frequency of XY males is 1 − *R*_*n*_, and the frequency of X’-sperm produced by X’Y males is *t* (while XY males produce X-sperm at a frequency of 0.5), the following equations are derived:

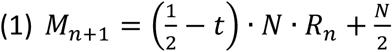

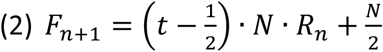

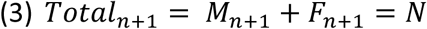

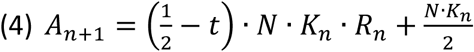

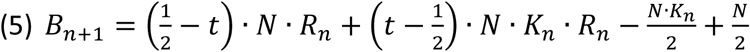

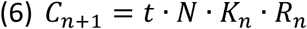

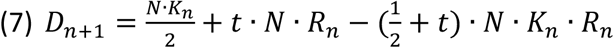

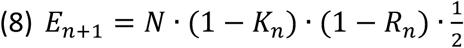

Thus, we can derive:

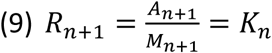

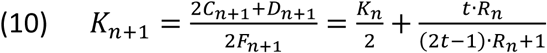

Thus,

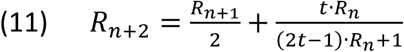

Equations (9), (10), and (11) define the recursive functions for *R*_*n*_ and *K*_*n*_, given the initial conditions *R*_0_ = *K*_0_ = 1/1000. These equations enable the calculation of *R*_*n*_ and *K*_*n*_ for any generation *n*. We found that as *n* increases, both *R*_*n*_ and *K*_*n*_ will eventually reach 1 and the rate depends on the value of *t*.

We further define the reproduction index *z* as the number of offspring a female produces, and the mating index *w* as the number of females a male can fertilize. The model requires *z* ≥ 2, because *z* = 1 leads to population decline even without a driver. Since *Drosophila* is a promiscuous species, we consider *w* ≥ 2 in our model. The parameter *w* is crucial in determining the population’s sensitivity to sex ratio distortion. In generation *n*, if *w* · *M*_*n*_ ≥ *F*_*n*_, there would be enough males to fertilize all females (all females are able to reproduce). Thus, the number of fertilized eggs produced in generation *n* is *N* = *z* · *F*_*n*_. In this scenario, the population will increase over generations, as each generation produces more females than the previous generation. The female-to-male ratio will also increase over generations. By incorporating *N* = *z* · *F*_*n*_ into equations (1) and (2), we get:

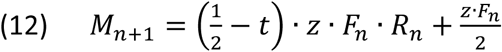

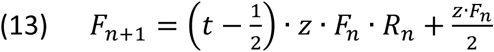

However, as the female-to-male ratio in the population continues to increase, eventually *w* ·*M*_*n*_ < *F*_*n*_, meaning some females will not be fertilized and thus will not reproduce. This will lead to population decline, as shown in Hamilton’s modeling (*9*). In this scenario, *w* · *M*_*n*_ females are able to reproduce, thus *N* = *z* · *w* · *M*_*n*_. By incorporating this into equations (1) and (2), we get:

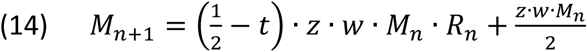

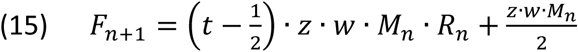

Using equations (12), (13), (14), and (15), combined with the recursive functions for *R*_*n*_ and *K*_*n*_ and the initial conditions (*M*_0_ = *F*_0_ = 1000), we can calculate *M*_*n*_, *F*_*n*_, and *Total*_*n*_ for each generation, when given specific values of parameters *t*, *z* and *w*. Population change rate *Total*_*n*+1_/*Total*_*n*_ and the male-to-female ratio *M*_*n*_/*F*_*n*_ can be calculated accordingly. Hamilton’s modeling (*9*) specifically addressed conditions where *t* = 1 (complete X chromosome drive), *z* = 2 (so that the population remains stable in the absence of driver), and *w* = 2. In this study, we explored how the population responds to a broader spectrum of *t*, *z* and *w* values.

The threshold drive strength can be calculated as follows. As *n* increases, if population reaches a steady state, then *M*_*n*+1_ = *M*_*n*_, *F*_*n*+1_ = *F*_*n*_, and *R*_*n*_ = *K*_*n*_ = 1. This steady state cannot occur when *w* · *M*_*n*_ ≥ *F*_*n*_, as the population would continue to increase as discussed above. We can mathematically prove this: if *w* · *M*_*n*_ ≥ *F*_*n*_, equation (13) yields *t*_*threshold*_ = 1/*z*. Since the model requires *z* ≥ 2, *t*_*threshold*_ = 1/*z* ≤ 0.5, which contradicts the premise that 0.5 < *t* ≤ 1. Therefore, a steady state is achievable only when *w* · *M*_*n*_ < *F*_*n*_. In this scenario, equation (14) yields 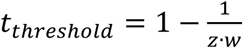 (fig. S8B). The minimum values of *z* and *w* (both equal to 2) yield the minimum *t*_*threshold*_ = 0.75 (fig. S8B).

---

**Fig. S1.**
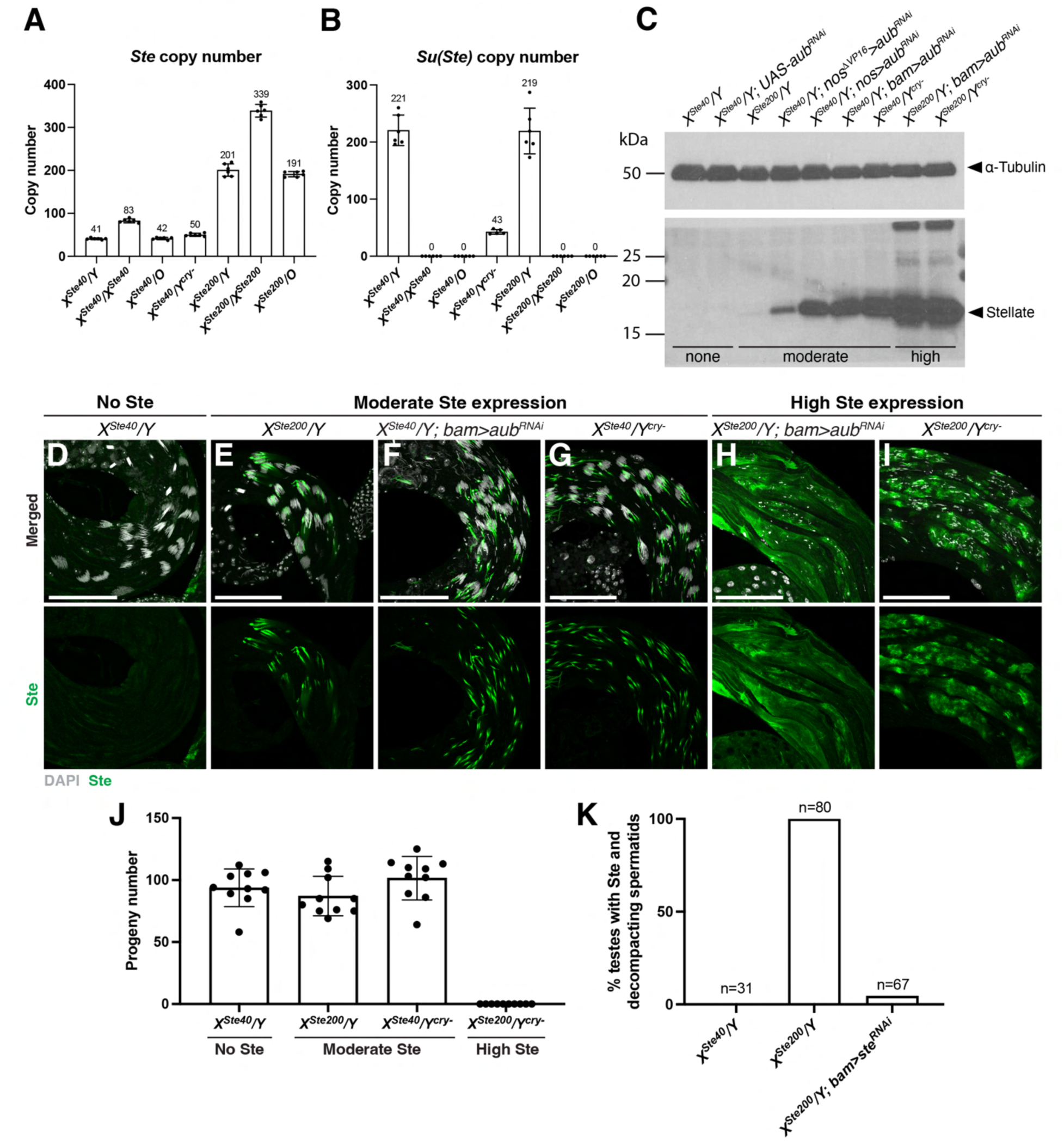
Varying degrees of *Ste* derepression in different genetic backgrounds. (**A**) *Ste* copy number on the X^Ste40^ and X^Ste200^ chromosomes, determined by ddPCR. *Ste* copy number in X/Y males is half of that in X/X females. The *Ste* copy number in X/Y males is the same as that in X/O males, demonstrating that the ddPCR primers and probes are specific to *Ste* and do not detect *Su(Ste)*. (**B**) *Su(Ste)* copy number on the normal Y and Y^cry-^ chromosomes, determined by ddPCR. *Su(Ste)* is not detected in X/O males or X/X females, demonstrating that the *Su(Ste)* ddPCR primers and probes are specific to *Su(Ste)* and do not cross-hybridize with *Ste*. (**C**) Western blotting to determine the expression level of Ste under the indicated genetic conditions, probed with anti-α-Tubulin (loading control) and anti-Ste antibodies. (**D** to **I**) Immunofluorescence staining for Ste (green) in testes from males of the indicated genotypes. Grey, DAPI. Scale bars, 100 μm. (**J**) Fertility assay for males of the indicated genotypes. Single males were crossed to two wild-type (*y w*) females for 5 days, and all progeny were counted. Ten males were assayed per genotype. (**K**) Percentage of testes expressing Ste and showing spermatid nuclear DNA compaction defects in the indicated genotypes. The number of testes scored for each genotype is shown above the bars.

**Fig. S2.**
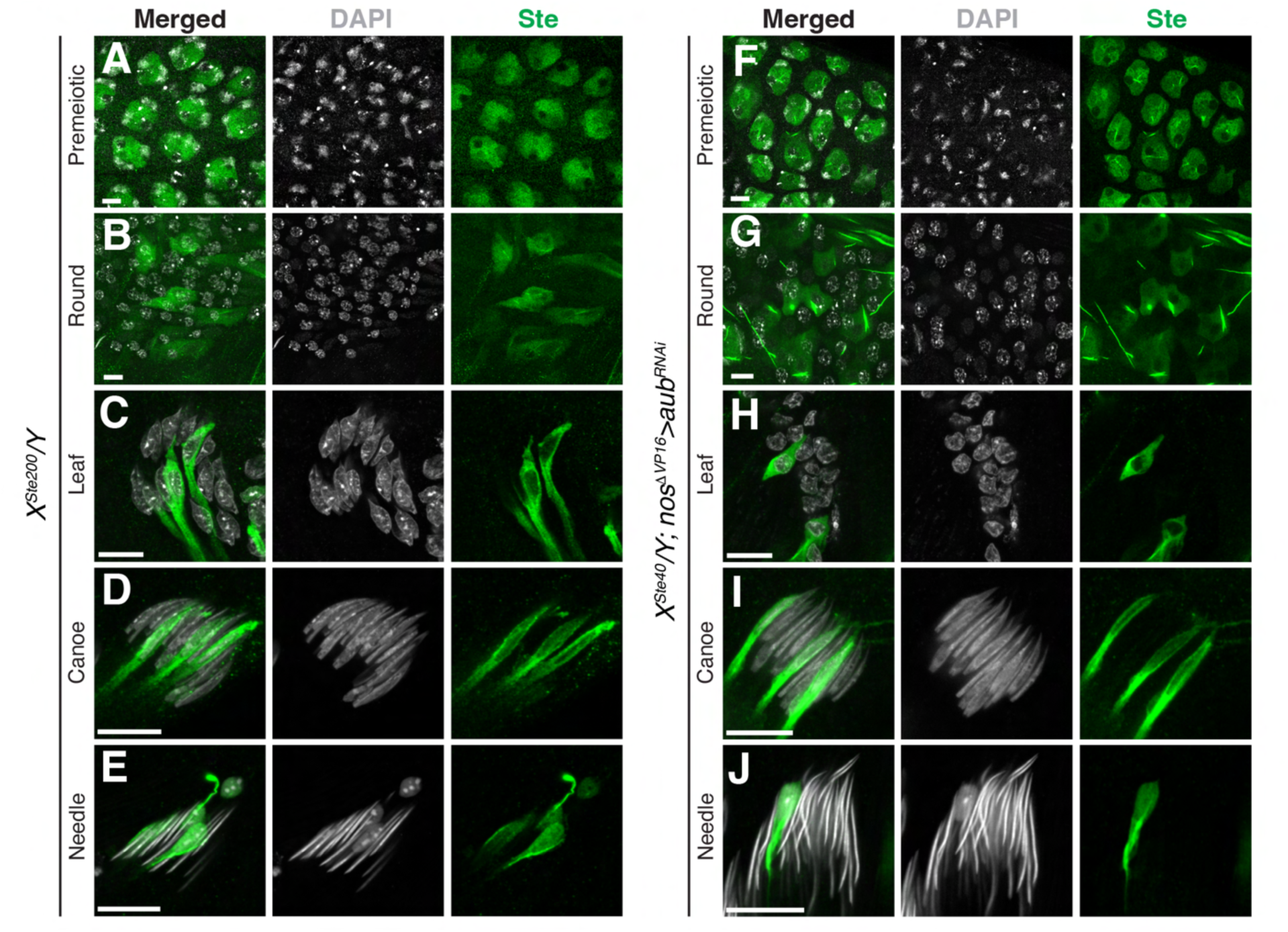
Moderately expressed Ste is localized to a subset of spermatids throughout spermiogenesis. (**A** to **E**) Immunofluorescence staining for Ste (green) in the indicated germ cell developmental stages in *X^Ste200^/Y* males: (A) Premeiotic spermatocytes; (B) round spermatids; leaf spermatids; (D) canoe spermatids; (E) needle spermatids. Grey, DAPI. Scale bars, 10 μm. (**F** to **J**) Immunofluorescence staining for Ste (green) in the indicated germ cell developmental stages in *X^Ste40^/Y; nos*^Δ*VP16*^*>aub^RNAi^* males: (F) Premeiotic spermatocytes; (G) round spermatids; (H) leaf spermatids; (I) canoe spermatids; (J) needle spermatids. Grey, DAPI. Scale bars, 10 μm.

**Fig. S3.**
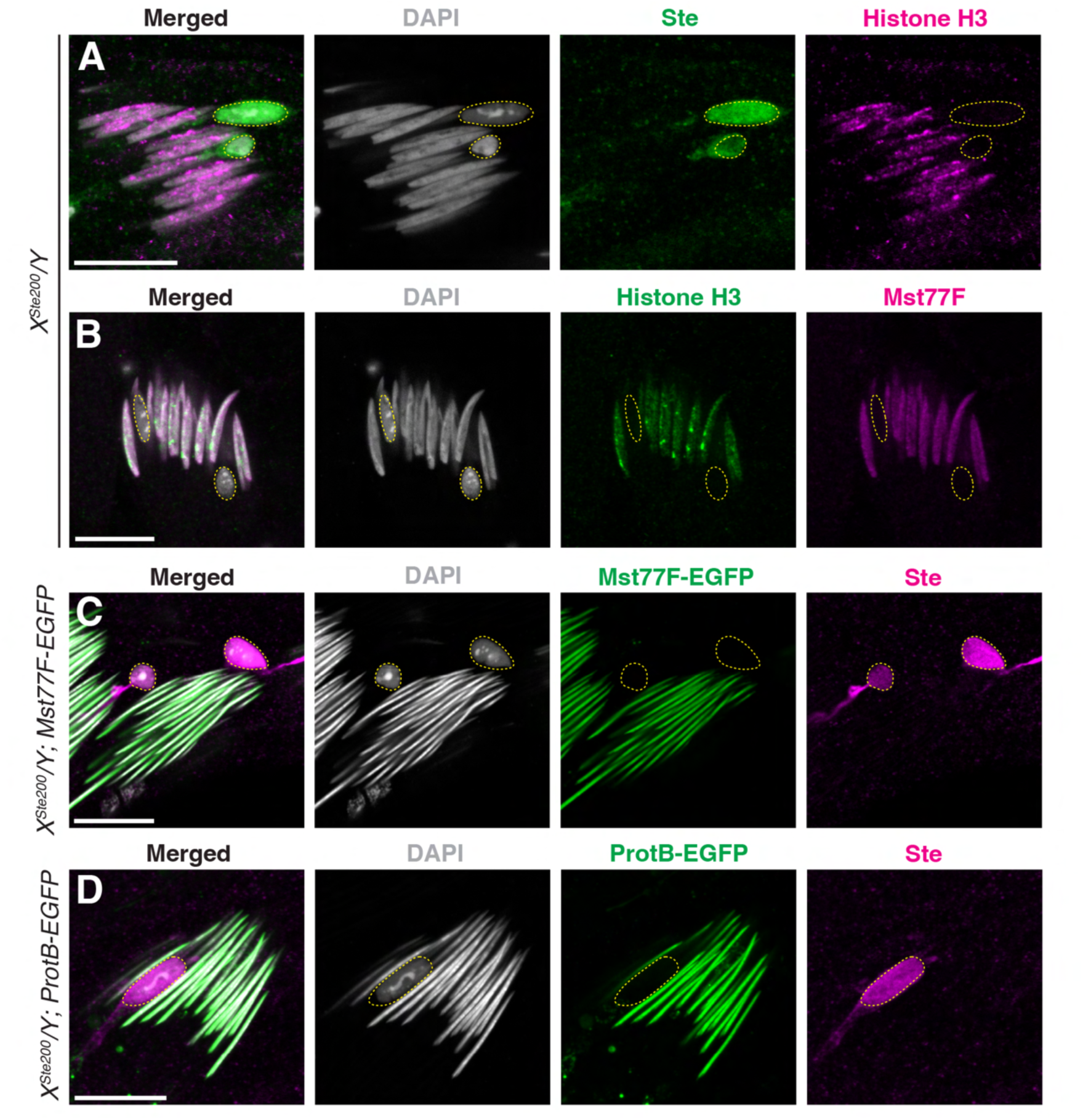
Ste interferes with the histone-to-protamine transition during spermiogenesis. (**A**) Canoe-stage spermatids from *X^Ste200^/Y* males stained for Ste (green), Histone H3 (magenta) and DAPI (grey), showing that Ste-containing spermatids (yellow dotted circles) do not have histones, whereas Ste-negative spermatids within the same cyst have Histone H3 signals. Scale bar: 10 μm. (**B**) Canoe-stage spermatids from *X^Ste200^/Y* males stained for Histone H3 (green), Mst77F (magenta, a protamine protein) and DAPI (grey), showing that spermatids that do not have Histone H3 and fail to condense DNA (presumably Ste-containing spermatids, yellow dotted circles) also lack Mst77F. Scale bar: 10 μm. (**C**) Needle-stage spermatids from *X^Ste200^/Y; Mst77F-EGFP* males (Mst77F-EGFP, green) stained for Ste (magenta) and DAPI (grey), showing that Ste-containing spermatids (yellow dotted circles) lack Mst77F. Scale bar, 10 μm. (C) Needle-stage spermatids from *X^Ste200^/Y; ProtamineB-EGFP* males (ProtamineB-EGFP, green) stained for Ste (magenta) and DAPI (grey), showing that Ste-containing spermatids (yellow dotted circle) lack Protamine B. Scale bar, 10 μm.

**Fig. S4.**
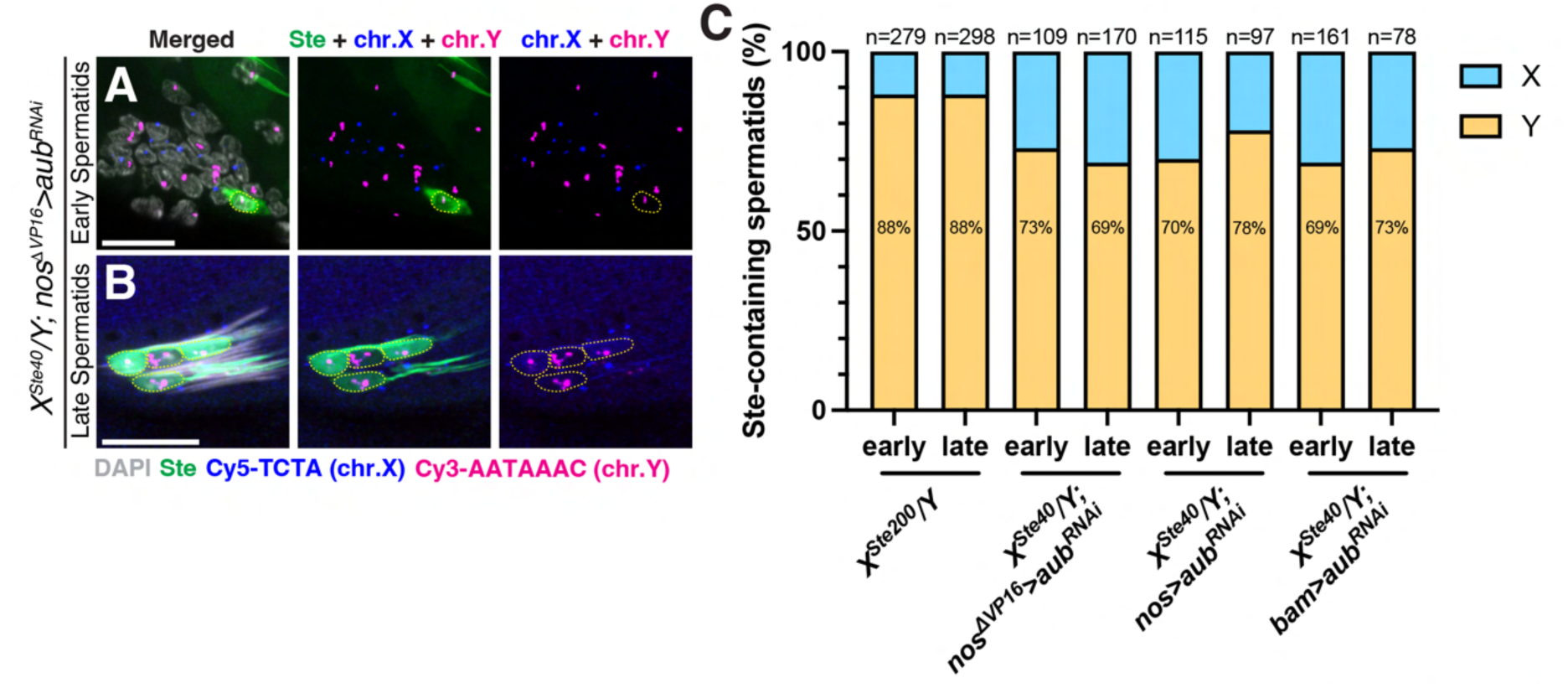
Ste is preferentially localized to Y-bearing spermatids throughout spermiogenesis. (**A** and **B**) Immunofluorescence and DNA-FISH of early (leaf stage) and late (needle stage) spermatids from *X^Ste40^/Y; nos*^Δ*VP16*^*>aub^RNAi^* males, stained for Ste (green), Cy5-TCTA (X chromosome, blue), and Cy3-AATAAAC (Y chromosome, magenta). Nuclei of Ste-containing spermatids are indicated by yellow dotted circles, showing that they predominantly contain Y chromosomes. Scale bars, 10 μm. (**C**) Frequency of X- or Y-bearing spermatids among Ste-containing spermatids in early (round + leaf) and late (canoe + needle) stages for the indicated genotypes.

**Fig. S5.**
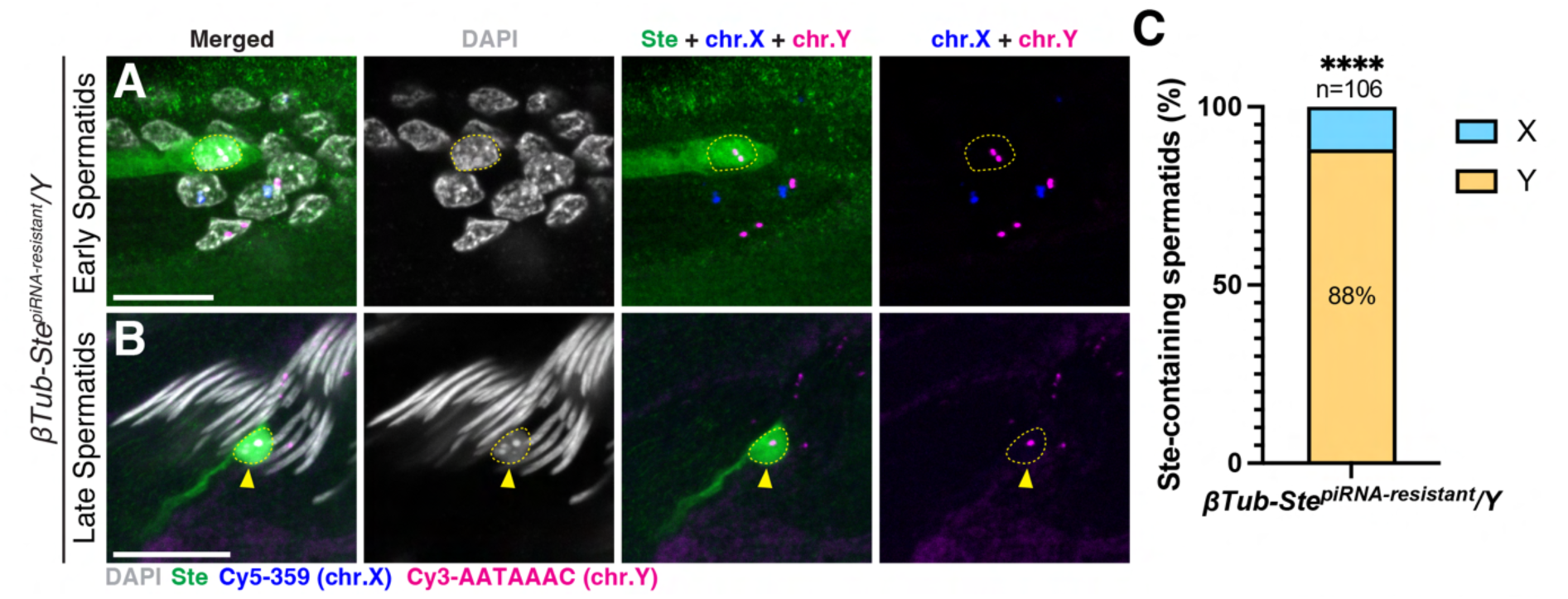
Ste is sufficient to cause Y-spermatid defects. (**A** and **B**) Immunofluorescence and DNA-FISH staining of early (leaf stage) (A) and late (needle stage) (B) spermatids from males expressing the piRNA-resistant *Ste* transgene (*βTub-Ste^piRNA-resistant^*), stained for Ste (green), X chromosome (Cy5-359, blue) and Y chromosome (Cy3-AATAAAC, magenta). Yellow dotted circles indicate the nuclei of Ste-containing spermatids, which predominantly contain a Y chromosome. Yellow arrowhead indicates a Ste-containing spermatid with nuclear DNA compaction defect. Grey, DAPI. Scale bars, 10 μm. (**C**) Percentage of X- or Y-bearing spermatids among Ste-containing spermatids in males expressing the piRNA-resistant *Ste* transgene (*βTub-Ste^piRNA-resistant^*). The number of scored Ste-containing spermatids is shown above the bar. Statistical analysis was performed using two-sided Fisher’s exact test (Null hypothesis: Ste-containing spermatids have equal chances of carrying X or Y chromosomes). \*\*\*\**P* < 0.0001.

**Fig. S6.**
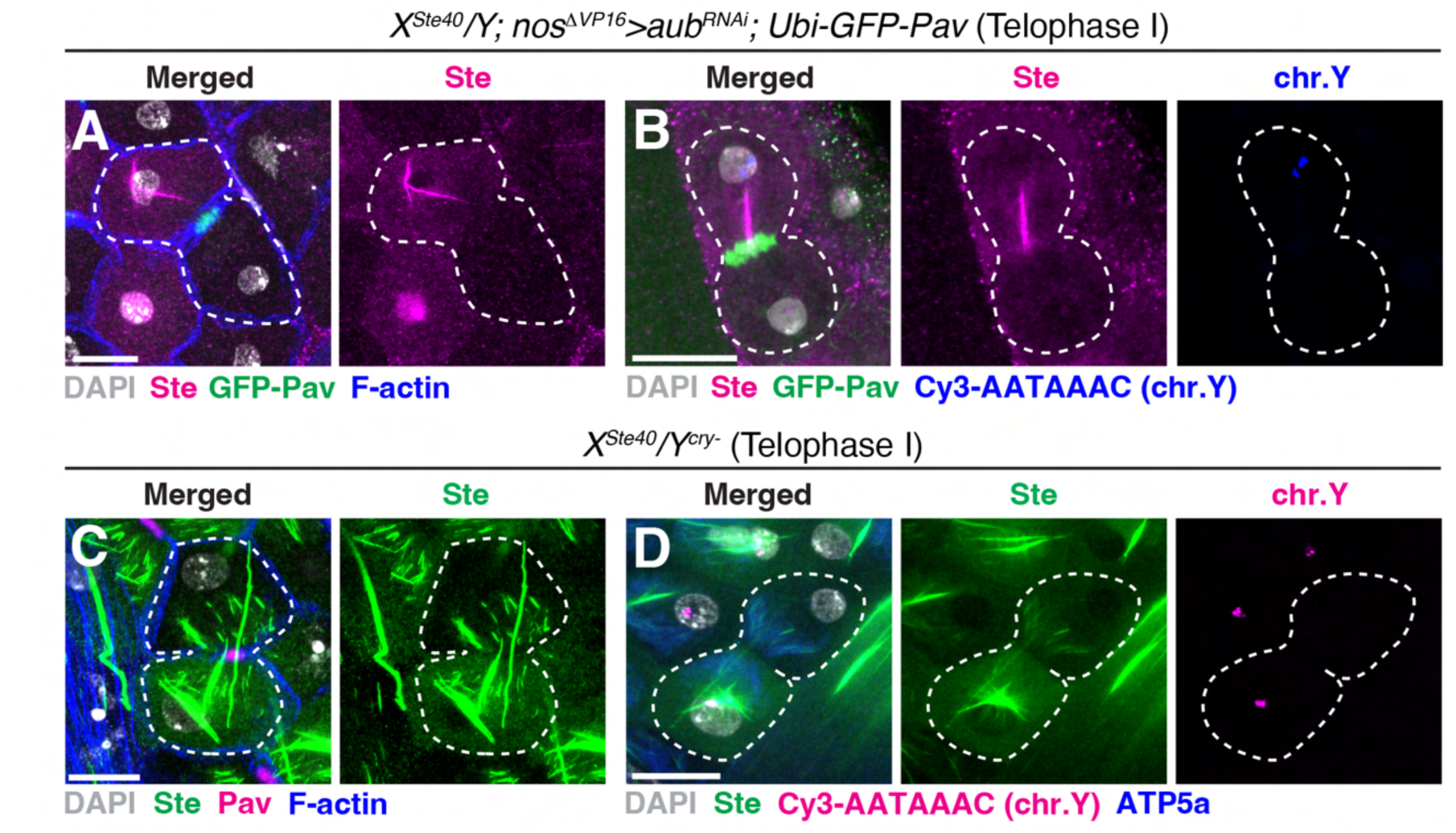
Ste asymmetrically segregates to Y chromosome-bearing cells during meiosis. **I.** (**A**) Immunofluorescence staining of a telophase I cell (indicated by white dotted lines) from *X^Ste40^/Y; nos*^Δ*VP16*^*>aub^RNAi^; Ubi-GFP-Pav* male, exhibiting asymmetric Ste segregation. Ste (magenta), GFP-Pav (green), and F-actin (blue). Grey, DAPI. Scale bar, 10 μm. (**B**) Immunofluorescence and DNA-FISH staining of a telophase I cell (indicated by white dotted lines) from *X^Ste40^/Y; nos*^Δ*VP16*^*>aub^RNAi^; Ubi-GFP-Pav* male, exhibiting asymmetric Ste segregation to the Y chromosome side. Ste (magenta), GFP-Pav (green), and the Y chromosome (Cy3-AATAAAC, blue). Grey, DAPI. Scale bar, 10 μm. (**C**) Immunofluorescence staining of a telophase I cell (indicated by white dotted lines) from *X^Ste40^/Y^cry-^* male, exhibiting asymmetric Ste segregation. Note that the relatively high expression of Ste in this genotype often resulted in both daughter cells inheriting Ste protein, although asymmetry remained apparent. Ste (green), Pav (magenta), and F-actin (blue). Grey, DAPI. Scale bar, 10 μm. (**D**) Immunofluorescence and DNA-FISH staining of a telophase I cell (indicated by white dotted lines) from *X^Ste40^/Y^cry-^* male, exhibiting asymmetric Ste segregation to the Y chromosome side. Ste (green), ATP5a (mitochondria decorating the spindle, blue), and the Y chromosome (Cy3-AATAAAC, magenta). Grey, DAPI. Scale bar, 10 μm.

**Fig. S7.**
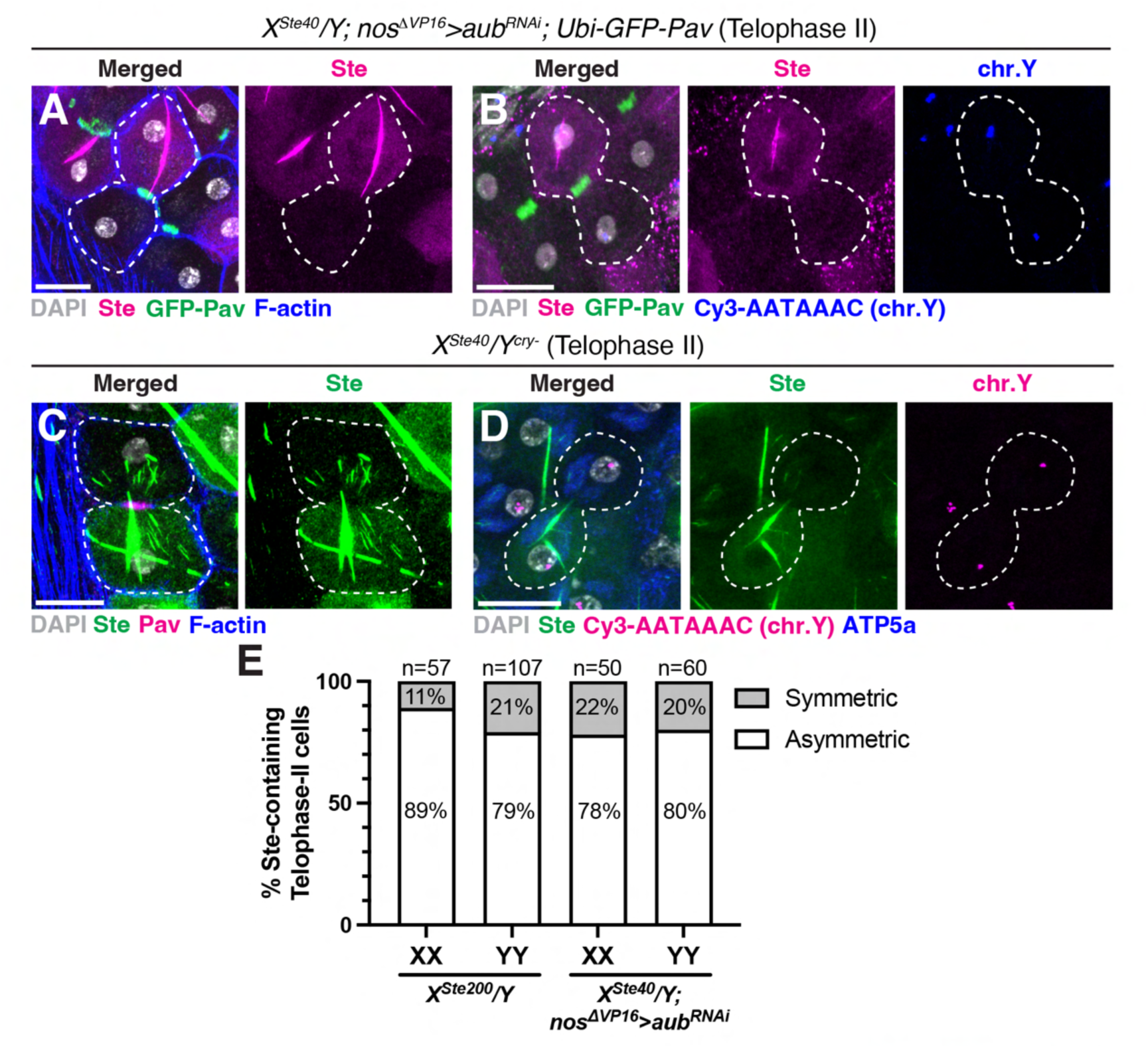
Ste segregates asymmetrically during meiosis II. (**A**) Immunofluorescence staining of a telophase II cell (indicated by white dotted lines) from *X^Ste40^/Y; nos*^Δ*VP16*^*>aub^RNAi^; Ubi-GFP-Pav* male, exhibiting asymmetric Ste segregation. Ste (magenta), GFP-Pav (green), and F-actin (blue). Grey, DAPI. Scale bar, 10 μm. (**B**) Immunofluorescence and DNA-FISH staining of a telophase II cell (indicated by white dotted lines) from *X^Ste40^/Y; nos*^Δ*VP16*^*>aub^RNAi^; Ubi-GFP-Pav* male. Ste (magenta), GFP-Pav (green), and the Y chromosome (Cy3-AATAAAC, blue). Grey, DAPI. Scale bar, 10 μm. (**C**) Immunofluorescence staining of a telophase II cell (indicated by white dotted lines) from *X^Ste40^/Y^cry-^* male, exhibiting asymmetric Ste segregation. Ste (green), Pav (magenta), and F-actin (blue). Grey, DAPI. Scale bar: 10 μm. (**D**) Immunofluorescence and DNA-FISH staining of a telophase II cell (indicated by white dotted lines) from *X^Ste40^/Y^cry-^* male, exhibiting asymmetric Ste segregation. Ste (green), ATP5a (mitochondria decorating the spindle, blue), and the Y chromosome (Cy3-AATAAAC, magenta). Grey, DAPI. Scale bar, 10 μm. (**E**) Frequency of Ste asymmetry during meiosis II among Ste-containing cells in the indicated genotypes. The number of scored meiosis II cells is shown above the bars.

**Fig. S8.**
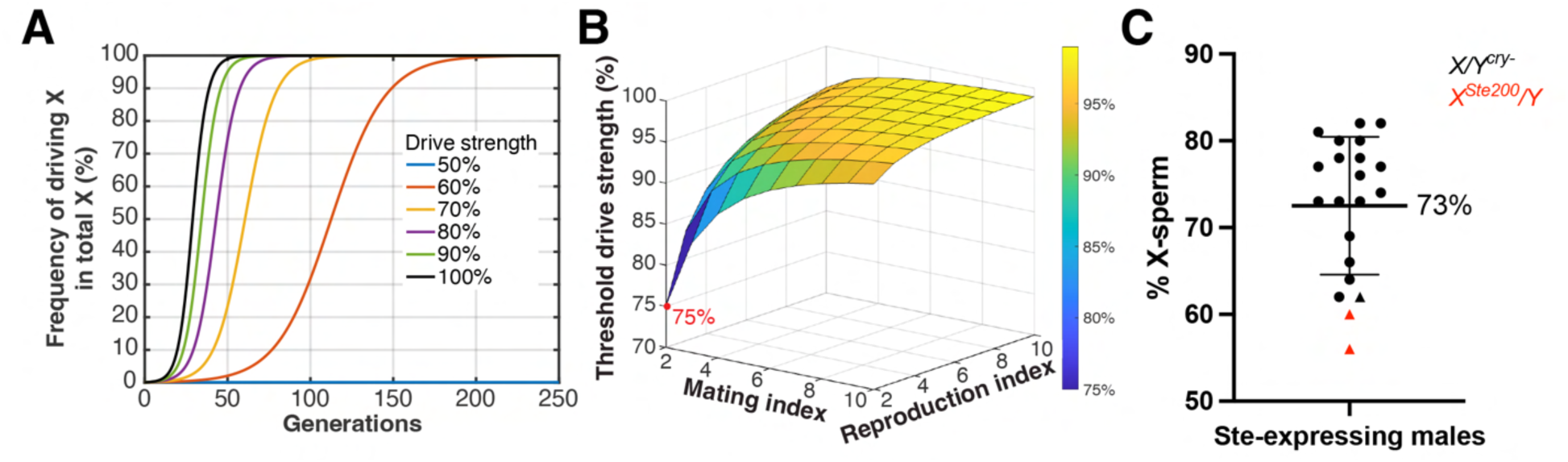
*Ste*’s drive strength is likely below the threshold drive strength. (**A**) The frequency of the driving X chromosome in the total X chromosome pool of the population over generations, plotted against varying degrees of drive strength. (**B**) Threshold drive strength with varying mating index and reproduction index. 75% is the lowest threshold drive strength (see Materials and Methods). (**C**) Percentage of X chromosome-bearing sperm produced by Ste-expressing males of the indicated genotypes. Each dot in the graph represents a genotype. Data include results from our own study (triangular dots) and previous studies (circular dots) (*16*). Mean ± SD is indicated in the graph. The number of scored progenies for each genotype is shown in table S2.

**Table S1.** Different levels of Ste expression lead to distinct phenotypes.

| Ste expression | Genotype | Phenotype |
| --- | --- | --- |
| None | $X^{Ste40}/Y$ | Wild type. |
| Moderate | $X^{Ste200}/Y$ | Ste localizes to a subset of spermatids, leading to the nuclear DNA compaction defect.<br>Males are fertile.<br>Males produce more females than males in the progeny. |
| | $X^{Ste40}/Y; nos^{\Delta VP16} > aub^{RNAi}$ | |
| | $X^{Ste40}/Y; nos > aub^{RNAi}$ | |
| | $X^{Ste40}/Y; bam > aub^{RNAi}$ | |
| | $X^{Ste40}/Y^{cry-}$ | |
| High | $X^{Ste200}/Y; bam > aub^{RNAi}$ | Ste causes the breakdown of entire spermatid cysts, resulting in no surviving sperm.<br>Males are sterile. |
| | $X^{Ste200}/Y^{cry-}$ | |

**Table S2.** Percentage of X-sperm produced by Ste-expressing males across various genotypes.

| Source | Genotype | Sperm genotype* |  | Sum | X:Y | X/(X+Y) |
| --- | --- | --- | --- | --- | --- | --- |
|  |  | X | Y |  |  |  |
| This study | *X<sup>Ste40</sup>/Y<sup>cry-</sup>* | 2924 | 1815 | 4739 | 1.6 | 62% |
|  | *X<sup>Ste200</sup>/Y; bam-gal4* | 2646 | 2044 | 4690 | 1.3 | 56% |
|  | *X<sup>Ste200</sup>/Y; Sp/CyO; TM2/TM6B* | 2475 | 1649 | 4124 | 1.5 | 60% |
| Palumbo et al., 1994 (16) | *W-12/Y<sup>cry-</sup>* | 1206 | 667 | 1873 | 1.8 | 64% |
|  | *Altamura-1/Y<sup>cry-</sup>* | 1567 | 966 | 2533 | 1.6 | 62% |
|  | *Fairfield-11/Y<sup>cry-</sup>* | 1261 | 354 | 1615 | 3.6 | 78% |
|  | *Salve-3/Y<sup>cry-</sup>* | 1369 | 705 | 2074 | 1.9 | 66% |
|  | *Salve-2/Y<sup>cry-</sup>* | 1720 | 374 | 2094 | 4.6 | 82% |
|  | *Altamura-46/Y<sup>cry-</sup>* | 1219 | 343 | 1562 | 3.6 | 78% |
|  | *Altamura-66/Y<sup>cry-</sup>* | 1645 | 576 | 2221 | 2.9 | 74% |
|  | *y w f/Y<sup>cry-</sup>* | 1186 | 529 | 1715 | 2.2 | 69% |
|  | *Altamura-40/Y<sup>cry-</sup>* | 1400 | 425 | 1825 | 3.3 | 77% |
|  | *Altamura-61/Y<sup>cry-</sup>* | 1506 | 447 | 1953 | 3.4 | 77% |
|  | *Valenzano-2/Y<sup>cry-</sup>* | 243 | 91 | 334 | 2.7 | 73% |
|  | *Sammichele/Y<sup>cry-</sup>* | 233 | 57 | 290 | 4.1 | 80% |
|  | *Giovinazzo/Y<sup>cry-</sup>* | 333 | 125 | 458 | 2.7 | 73% |
|  | *Altamura-4/Y<sup>cry-</sup>* | 467 | 175 | 642 | 2.7 | 73% |
|  | *Altamura-22/Y<sup>cry-</sup>* | 373 | 94 | 467 | 4.0 | 80% |
|  | *Salve-4/Y<sup>cry-</sup>* | 120 | 29 | 149 | 4.1 | 81% |
|  | *Altamura-36/Y<sup>cry-</sup>* | 184 | 41 | 225 | 4.5 | 82% |
|  | *Gandoli-6/Y<sup>cry-</sup>* | 183 | 58 | 241 | 3.2 | 76% |
| Average | --- |  |  |  | 2.9 | 72.5% |

\* Non-disjunction products (XY- and O-sperm) are excluded in our analysis.

**Table S3.**
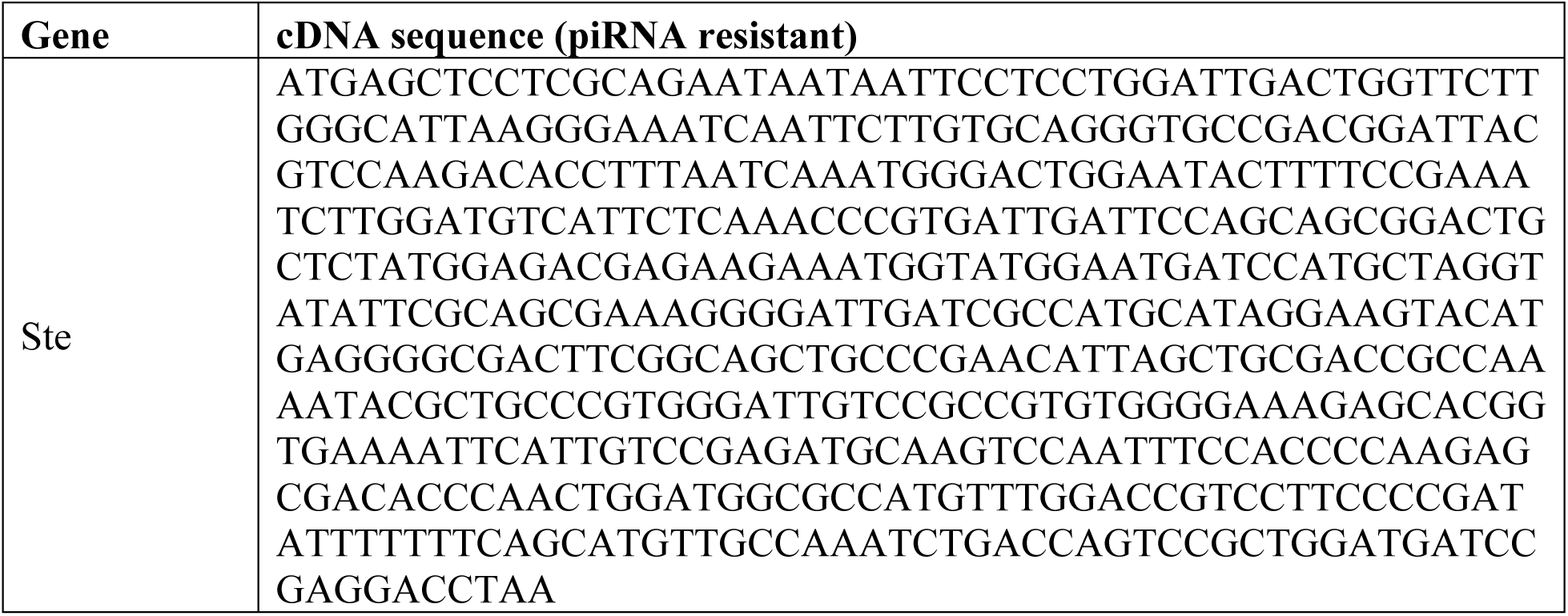
cDNA sequence of piRNA-resistant *Ste*.

**Table S4.** Fertility assay for males expressing different levels of Ste.

| **Ste expression** | **No** | **Moderate** |  | **High** |
| --- | --- | --- | --- | --- |
| **Genotype** | $X^{Ste40}/Y$ | $X^{Ste200}/Y$ | $X^{Ste40}/Y^{cry-}$ | $X^{Ste200}/Y^{cry-}$ |
| male 1 | 105 | 75 | 125 | 0 |
| male 2 | 89 | 80 | 113 | 0 |
| male 3 | 92 | 76 | 110 | 0 |
| male 4 | 94 | 115 | 64 | 0 |
| male 5 | 103 | 85 | 89 | 0 |
| male 6 | 106 | 102 | 109 | 0 |
| male 7 | 93 | 75 | 86 | 0 |
| male 8 | 85 | 85 | 114 | 0 |
| male 9 | 58 | 69 | 103 | 0 |
| male 10 | 112 | 109 | 102 | 0 |

**Table S5.** The distorted sex ratio in Ste-expressing males is rescued by Ste RNAi.

| **Genotype** | ***X<sup>Ste40</sup>/Y; bam-gal4*** |  |  | ***X<sup>Ste200</sup>/Y; bam-gal4*** |  |  | ***X<sup>Ste200</sup>/Y; bam>ste<sup>RNAi</sup>*** |  |  |
| --- | --- | --- | --- | --- | --- | --- | --- | --- | --- |
| **Individual** | **Female** | **Male** | **Female/Male** | **Female** | **Male** | **Female/Male** | **Female** | **Male** | **Female/Male** |
| male 1 | 284 | 264 | 1.08 | 301 | 220 | 1.37 | 282 | 243 | 1.16 |
| male 2 | 216 | 221 | 0.98 | 281 | 216 | 1.30 | 260 | 244 | 1.07 |
| male 3 | 278 | 277 | 1.00 | 289 | 260 | 1.11 | 292 | 240 | 1.22 |
| male 4 | 254 | 265 | 0.96 | 338 | 227 | 1.49 | 302 | 258 | 1.17 |
| male 5 | 249 | 226 | 1.10 | 264 | 238 | 1.11 | 269 | 282 | 0.95 |
| male 6 | 222 | 295 | 0.75 | 283 | 262 | 1.08 | 288 | 253 | 1.14 |
| male 7 | 285 | 265 | 1.08 | 196 | 121 | 1.62 | 173 | 169 | 1.02 |
| male 8 | 164 | 192 | 0.85 | 185 | 167 | 1.11 | 160 | 161 | 0.99 |
| male 9 | 111 | 119 | 0.93 | 150 | 104 | 1.44 | 206 | 188 | 1.10 |
| male 10 | 242 | 273 | 0.89 | 359 | 229 | 1.57 | 296 | 239 | 1.24 |

**Table S6.** Quantification of Ste-containing spermatids.

| Genotype | Ste-containing spermatids |  |  |  | Y/X+Y (%) | Non-disjunction (%) |
| --- | --- | --- | --- | --- | --- | --- |
|  | X | Y | XY or O | Total |  |  |
| *X<sup>Ste200</sup>/Y* | 70 | 507 | 13 | 590 | 88% | 2% |
| *X<sup>Ste40</sup>/Y; nos<sup>DVP16</sup>>aub<sup>RNAi</sup>* | 82 | 197 | 11 | 290 | 71% | 4% |
| *X<sup>Ste40</sup>/Y; nos>aub<sup>RNAi</sup>* | 56 | 156 | 1 | 213 | 74% | 0% |
| *X<sup>Ste40</sup>/Y; bam>aub<sup>RNAi</sup>* | 71 | 168 | 6 | 245 | 70% | 2% |
| *βTub-Ste<sup>piRNA-resistant</sup>/Y* | 13 | 93 | 0 | 106 | 88% | 0% |

**Table S7.** Fisher’s exact test results for Ste’s preferential localization in spermatids.

| Genotype |  | Ste-containing spermatids |  |  | P value | Asterisks |
| --- | --- | --- | --- | --- | --- | --- |
|  |  | X | Y | Total |  |  |
| *X<sup>Ste200</sup>/Y* | Expected | 288 | 289 | 577 | <0.0001 | **** |
|  | Observed | 70 | 507 | 577 |  |  |
| *X<sup>Ste40</sup>/Y; nos<sup>DVP16</sup>>aub<sup>RNAi</sup>* | Expected | 139 | 140 | 279 | <0.0001 | **** |
|  | Observed | 82 | 197 | 279 |  |  |
| *X<sup>Ste40</sup>/Y; nos>aub<sup>RNAi</sup>* | Expected | 106 | 106 | 212 | <0.0001 | **** |
|  | Observed | 56 | 156 | 212 |  |  |
| *X<sup>Ste40</sup>/Y; bam>aub<sup>RNAi</sup>* | Expected | 119 | 120 | 239 | <0.0001 | **** |
|  | Observed | 71 | 168 | 239 |  |  |
| *bTub-Ste<sup>piRNA-resistant</sup>/Y* | Expected | 53 | 53 | 106 | <0.0001 | **** |
|  | Observed | 13 | 93 | 106 |  |  |

Null hypothesis: Ste-containing spermatids have equal probabilities of containing either X or Y chromosomes.

**Table S8.**
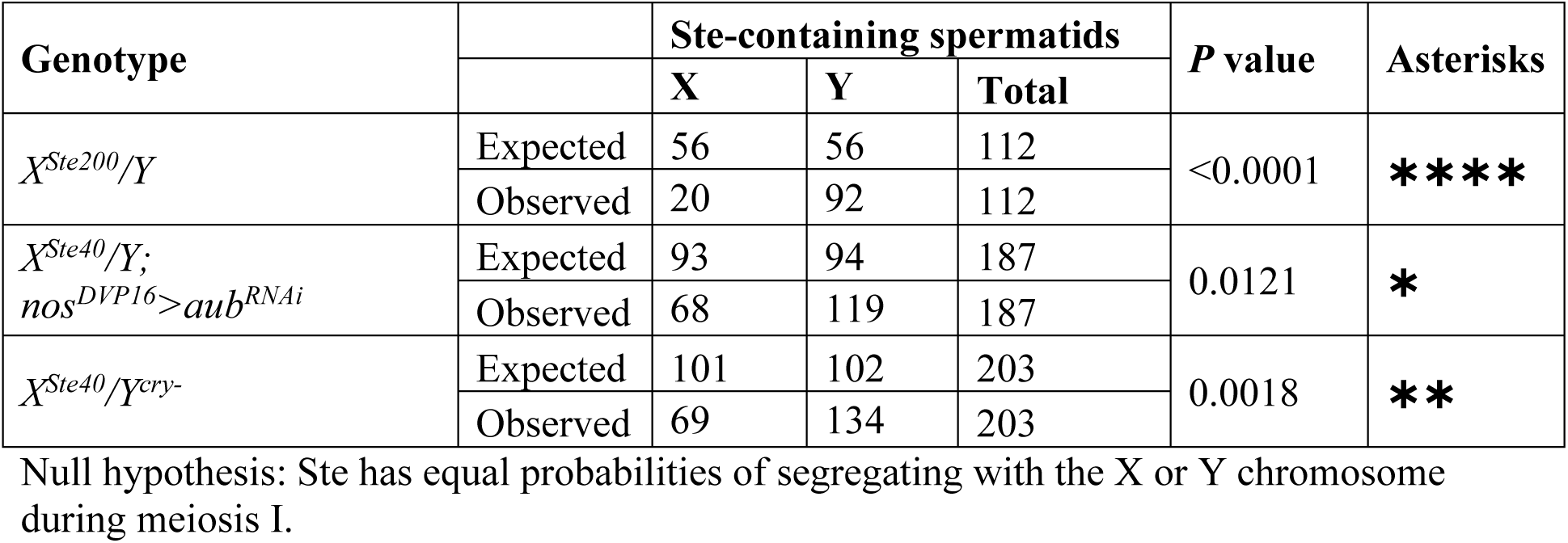
Fisher’s exact test results for the association of Ste with the Y chromosome in meiosis I.

## Notes

### Competing Interest Statement

The authors have declared no competing interest.

